# Super-resolution compatible DNA labeling technique reveals chromatin mobility and organization changes during differentiation

**DOI:** 10.1101/2025.02.06.636912

**Authors:** Maruthi K. Pabba, Miroslav Kuba, Tomáš Kraus, Kerem Celikay, Janis Meyer, Sunik Kumar Pradhan, Andreas Maiser, Hartmann Harz, Heinrich Leonhardt, Karl Rohr, Michal Hocek, M. Cristina Cardoso

**Affiliations:** Cell Biology and Epigenetics, Department of Biology, Technical University of Darmstadt, 64287 Darmstadt, Germany; Institute of Organic Chemistry and Biochemistry, Czech Academy of Sciences, CZ-16000 Prague 6, Czech Republic; Department of Organic Chemistry, Faculty of Science, Charles University, CZ-12843 Prague 2, Czech Republic; Biomedical Computer Vision Group, BioQuant, IPMB, Heidelberg University, Germany; Human Biology and Bioimaging, Faculty of Biology, Ludwig Maximilians University Munich, 81377 Munich, Germany

**Keywords:** dC^SiR^TP, SNTT1, chromatin mobility, neural differentiation, STED, human iPSC

## Abstract

Chromatin dynamics play a crucial role in cellular differentiation, yet tools for studying global chromatin mobility in living cells remain limited. Here, we developed a novel probe for the metabolic labeling of chromatin and tracking its mobility during neural differentiation. The labeling system utilizes a newly developed silicon rhodamine-conjugated deoxycytidine triphosphate (dC^SiR^TP). We show that this dCTP is efficiently delivered into living human induced pluripotent stem cells (iPSCs) and neural stem cells (NSCs) via a synthetic transporter (SNTT1). Using correlative confocal microscopy and stimulated emission depletion (STED) super-resolution microscopy, we quantified the sizes of labeled chromatin domains. Time lapse super-resolution microscopy combined with single particle tracking revealed that chromatin mobility decreases during the transition from iPSCs (pluripotent state) to NSCs and neurons (differentiated state). This reduction in mobility correlates with the differentiation state, suggesting a role for chromatin dynamics in cellular plasticity. Concomitant mechanistic insights obtained from MNase digestion assays, chromatin compaction and histone modification analyses revealed a decrease in chromatin accessibility during neuronal differentiation, indicating that chromatin adopts a more constrained and compacted structure. These findings provide new insights into chromatin regulation during neurogenesis.

## INTRODUCTION

The three-dimensional organization and dynamics of chromatin play crucial roles in regulating gene expression and cellular identity (1) (2). While the structural aspects of chromatin organization have been extensively studied through techniques like Hi-C and microscopy, our understanding of real-time chromatin mobility during cellular differentiation remains limited (3) (4). This dynamic behavior of chromatin is particularly relevant in the context of cellular differentiation, where large-scale reorganization of the genome occurs to establish and maintain cell type-specific gene expression programs (5).

The differentiation of human induced pluripotent stem cells (iPSCs) into neurons represents an excellent model system to study chromatin dynamics during cell fate transitions. Pluripotent stem cells are characterized by a unique nuclear architecture, featuring a more open and accessible chromatin landscape that is thought to contribute to their developmental plasticity (6) (7). During neural differentiation, dramatic changes in chromatin architecture have been documented. These include the formation of new topologically associating domains (TADs), the establishment of neural-specific enhancer-promoter interactions and the separation of neuronal lineage specific genes from the nuclear lamina (8) (9).

Recent single-cell studies have revealed unprecedented heterogeneity in chromatin states during neural differentiation, with distinct chromatin accessibility patterns emerging even before fate-determining transcriptional programs are activated (10). Advanced imaging techniques have shown that chromatin compaction states correlate strongly with cell fate commitment, where regions containing neural-specific genes undergo significant structural reorganization during differentiation (11). Notably, disruption of chromatin remodeling factors during neural differentiation can lead to severe neurodevelopmental disorders, highlighting the critical importance of proper chromatin dynamics in neural development (12).

Recent advances in fluorescent probes and super-resolution microscopy have opened new possibilities for studying chromatin dynamics in living cells. The development of synthetic nucleoside triphosphate transporters (SNTTs) capable of delivering modified nucleotides has provided novel tools for labeling chromatin while maintaining cellular viability (13). Several types of modified dNTPs have been successfully used in combination with SNTT for staining and metabolic labeling of genomic DNA in cellulo. These include nucleotides bearing bright fluorescent (14) or environment-sensitive fluorophores (15) (16) or reactive groups for post synthetic fluorogenic click reactions (17). On the other hand, some other fluorescent dNTPs have not shown in cellulo incorporation to genomic DNA (18) or exerted high toxicity (19), which prevented their applications. However, none of these fluorophores or probes were suitable for advanced imaging techniques such as stimulated emission depletion (STED) microscopy, which offers superior spatial resolution for imaging chromatin. Recent developments in live-cell imaging have enabled visualization of chromatin dynamics with unprecedented temporal resolution, revealing rapid local reorganization events that occur on timescales of seconds to minutes (20) (21, 22).

Here, we present a novel approach to study global chromatin structure and dynamics throughout neural differentiation in human cells using a newly developed dC^SiR^TP nucleotide analog compatible with STED microscopy, combined with the synthetic nucleoside triphosphate transporters (SNTT1). This allowed us to directly label and measure chromatin structures in living human cells at different spatial resolutions and to track the dynamics of these structures throughout neural differentiation.

## RESULTS & DISCUSSION

### Synthesis of STED-compatible fluorescently labeled nucleotides and labeling of chromatin

Silicon rhodamine-modified nucleoside triphosphate dC^SiR^TP was prepared by strain-promoted azide-alkyne cycloaddition reaction of azido-propargyloxy-triethylene glycol-conjugated deoxycytidine triphosphate dC^pegN3^TP (16) with bicyclononyne-linked silicon rhodamine fluorophore SiR-BCN (Figure 1A). The product was isolated in 50% yield after purification by reverse-phase high-performance liquid chromatography (HPLC). The solution of modified nucleotide dC^SiR^TP in phosphate buffered saline (PBS) showed an absorption maximum and an emission maximum at 651 nm and 672 nm, respectively (Table S1 and Figure S1A). The corresponding extinction coefficient and quantum yield in PBS were calculated to be 124800 M^-1^cm^-1^ and 54%, respectively.

**Figure 1.**
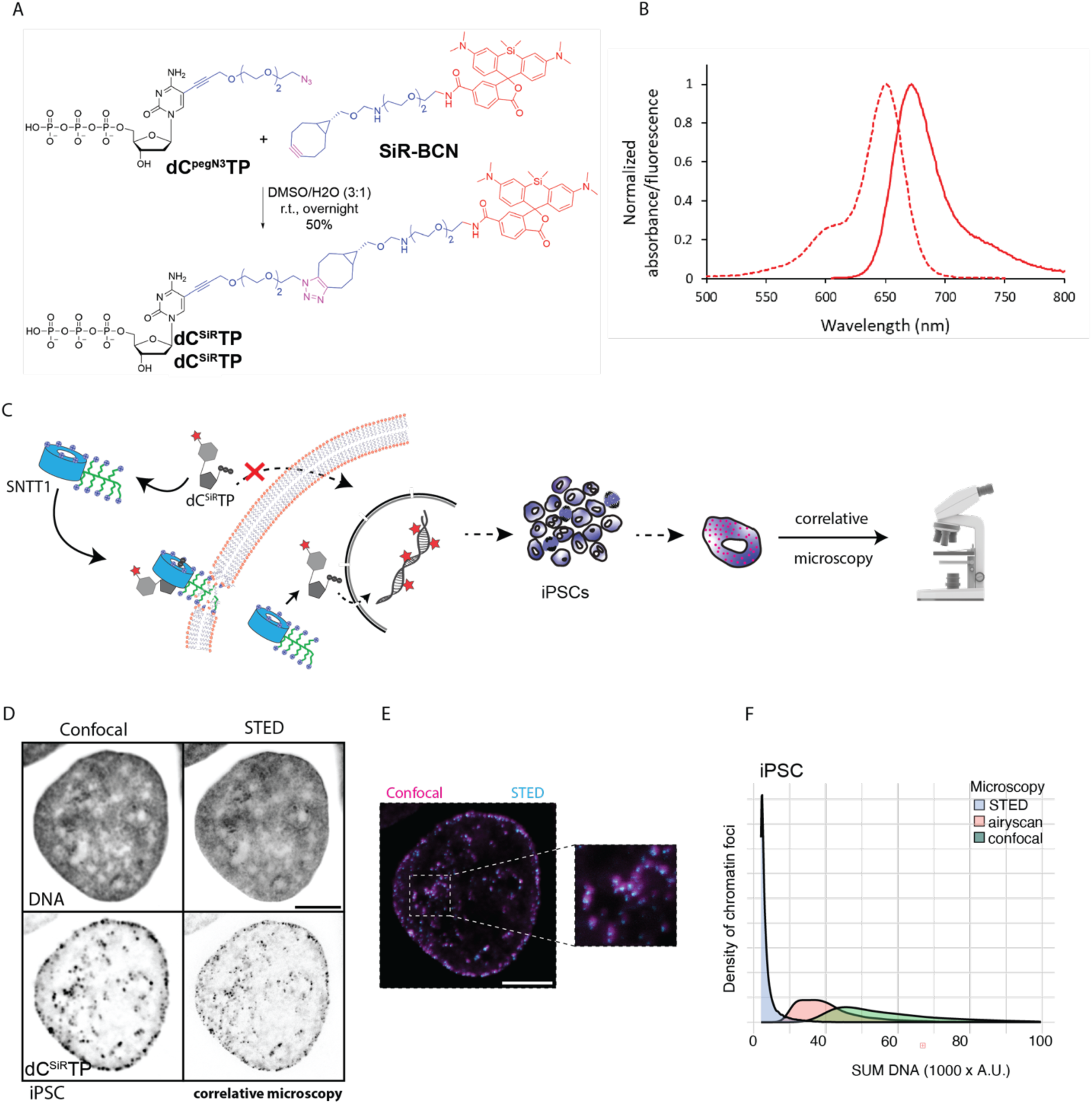
Synthesis and chromatin labeling using dC^SiR^TP. **(A)** Chemical synthesis and metabolic DNA labeling in living cells. SiR-BCN click reaction with azido-terminated dC^pegN3^TP precursor. **(B)** Absorption and fluorescence spectrum of DNA19_1C^SiR^ in PBS. **(C)** A schematic representation of transport of dC^SiR^TP across the cell membrane by synthetic nucleoside triphosphate transporter (SNTT1) and subsequent visualization of labeled DNA foci by correlative microscopy. **(D-F)** Chromatin labeled with dC^SiR^TP and imaged using correlative microscopy including stimulated emission depletion (STED) microscopy, confocal microscopy and Airyscan joint deconvolution microscopy (Table S6). The representative images show the total DNA and dC^SiR^TP labeled chromatin sites in grey scale and overlay image of dC^SiR^TP foci in cyan - STED, magenta - confocal. The SUM DNA (total DNA content) was plotted for different spatial resolutions. Scale bars: 5 µm.

The modified dC^SiR^TP was tested as a substrate for KOD XL DNA polymerase in primer extension (PEX) and polymerase chain reaction (PCR) experiments. PEX with 19-mer (Table S2) encoding for incorporation of a single modification proceeded smoothly and the product was detected by polyacrylamide gel electrophoresis (PAGE) (Figure S2A) and by MALDI-TOF mass spectrometry analysis (Figure S7). The SiR-labeled DNA in PBS showed and absorption maximum and an emission maximum at 651 nm and 672 nm, respectively (Figure 1B), with a quantum yield of 48% (Table S1). Thus, no significant change in the photophysical properties of SiR fluorophore was observed. PCR was tested using mixtures with varying ratios of dC^SiR^TP and dCTP (0-100%). Amplicon formation was detectable with up to 60% of dC^SiR^TP used in the reaction mixture. (Figure S2B, see Table S2, S3 for primer and template sequences).

Using the modified dC^SiR^TP nucleotide, with a STED-compatible (super resolution microscopy) fluorophore, we proceeded to test it on cultures of human induced pluripotent stem cells (iPSCs) (Table S4). Firstly, the iPSCs were checked for pluripotency using immunofluorescence detection of the pluripotent markers Sox2 and Oct 3/4 and active DNA replication was verified using incorporation and detection of EdU. The iPSC cultures were positive for pluripotent markers and were actively replicating the genomic DNA (Figure S8).

Subsequently, we labeled chromatin by incubating cells in a tricine buffer (pH 7.4) containing 10 µM SNTT1 and 10 µM dC^SiR^TP for 15 minutes at 37 °C (Table S5). SNTT1 facilitated the transport of labeled nucleotides into the cells allowing for an efficient incorporation of the labeled nucleotides into the genome of the replicating cells. This labeling process, which depends on active DNA replication, allows for a broad, random distribution of labeled genomic regions, providing unbiased coverage of the entire genome. Rapid chromatin labeling (within 10–15 minutes) was achieved without disrupting cell morphology by avoiding the trypsinization required by electroporation-based intracellular delivery methods. This allowed immediate imaging (Movie 1). After labeling chromatin in live cells with dC^SiR^TP and staining of total DNA with a live cell DNA dye, the iPSCs were fixed for correlative microscopic imaging.

Cells with labeled chromatin (dC^SiR^TP) were imaged comparing different spatial resolution techniques including STED microscopy, laser scanning confocal microscopy, and Airy scan microscopy (Figure 1C, Table S6). Representative gray-scale images of the DNA dye (Abberior 590 DNA dye) and dC^SiR^TP obtained from both confocal and STED imaging are shown in Figure 1D. The overlay image (Figure 1E) displays the same labeled chromatin visualized with STED (cyan) and confocal (magenta) microscopy. We observed that each clustered focus detected by confocal microscopy contained multiple, distinct foci when resolved with STED microscopy (Figure 1E). We then segmented the labeled foci at different spatial resolutions and plotted (Figure 1F) the foci count (y-axis: density) against the total DNA content within the segmented foci (x-axis: SUM DNA). The results demonstrated that STED microscopy provided significantly higher resolution of chromatin structure compared to confocal microscopy and Airy scan imaging.

### Chromatin mobility decreases upon neural differentiation

We next performed in vitro differentiation of iPSCs to obtain precursor neural stem cells (NSCs) as described (supplementary methods 6, Table S4, Table S7, Figure S9). Then, we established a protocol for differentiation of NSCs to neurons (supplementary methods 6, Table S7). Differential interference contrast (DIC) images of live cells, where NSCs were differentiated into neurons for 10 days are shown in Figure S10. The DIC images clearly show the neuronal network showing successful neuronal differentiation (Figure S10). To further visualize the cellular morphology of iPSCs, NSCs, and neurons, we acquired DIC images concomitantly with images of cells stained with the live-cell DNA dye (cyan) and a tubulin live-cell dye (red) (Figure 2A). Clear morphological differences between the differentiated stages were evident with the double live-cell dye staining.

**Figure 2.**
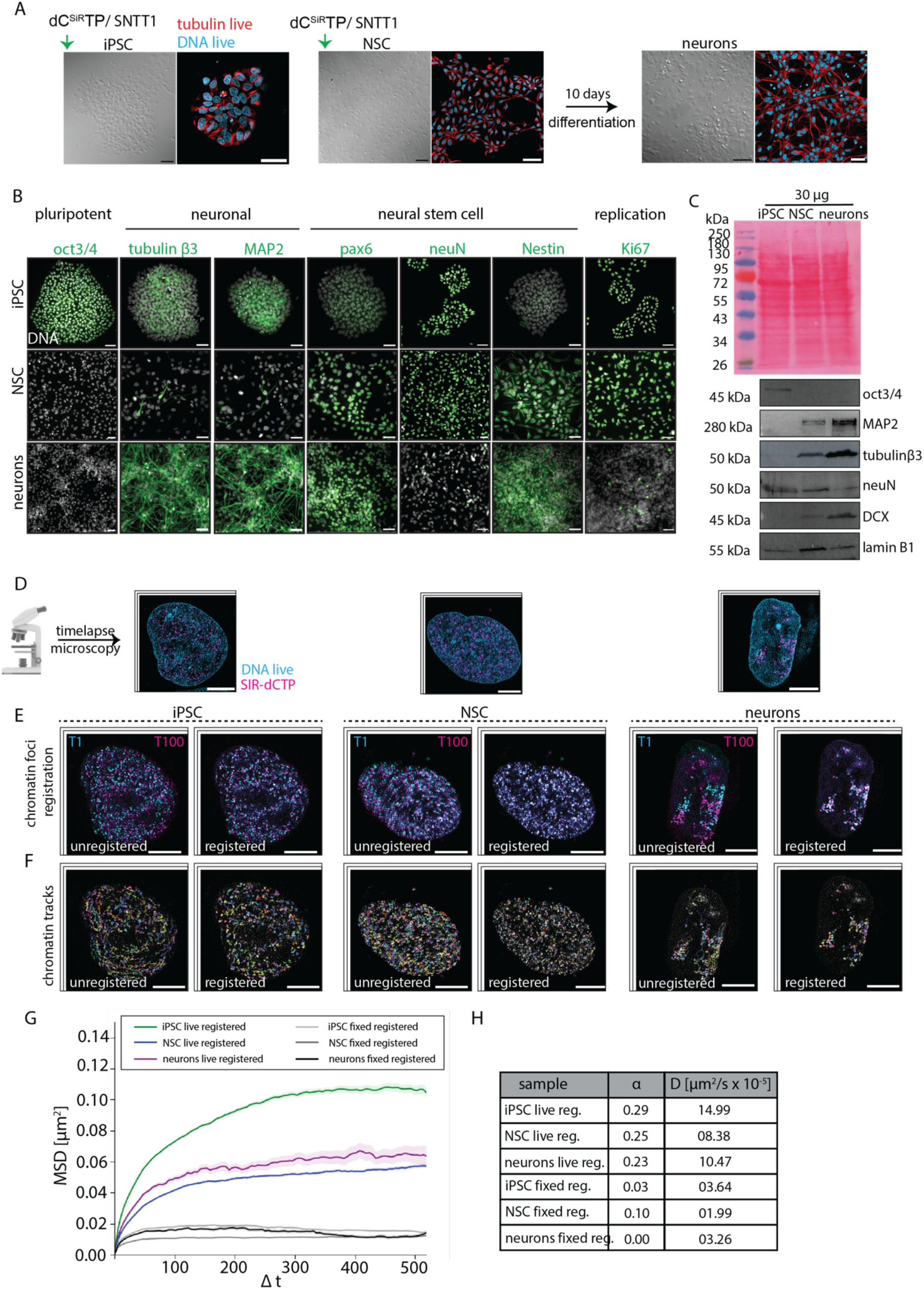
Characterization and chromatin mobility changes during neuronal differentiation. **(A)** Differential interference contrast (DIC) and tubulin (red) + DNA (cyan) live cell dye images for iPSC, NSC and neurons. Scale bars: 50 µm (black) and 10 µm (white). **(B)** Immunodetection of pluripotent and neuronal lineage specific markers in iPSC, NSC and neurons (Methods, Table S8). Scale bars: 50 µm. **(C)** Western blot detection of pluripotent and neuronal lineage specific markers in iPSC, NSC and neurons (2 biological replicates). **(D)** Chromatin labeled iPSC, NSC and neurons using SNTT1 + dC^SiR^TP complex (DNA - cyan, dC^SiR^TP - magenta) (> 4 biological replicates). **(E)** Labeled chromatin in all cell types was imaged using live cell microscopy and foci from time frame 1 (T1-cyan) was overlaid with time frame 100 (T100-magenta). Scale bars: 10 µm. **(F)** Registration of chromatin was performed using the live cell DNA dye as reference to estimate the actual chromatin mobility. The tracks in unregistered versus registered movies over time are illustrated to show the importance of 3D live cell registration. **(G)** Mean square displacement (y-axis, µm^2^) over time (x-axis, Δt) for iPSC (green), NSC (blue) and neurons (magenta). The fixed cells MSD (gray shade) for each cell differentiation state was plotted in grey. **(H)** The diffusion constant (D - µm^2^/s x 10^-5^) and ɑ values are listed.

To validate the in vitro differentiation of iPSCs into neurons, we performed immunofluorescence staining on iPSCs, NSCs, and neurons using specific markers for pluripotency, neural stem cells, and mature neurons (Figure 2B, Figure S11, Figure S12). Immunostaining revealed that only iPSCs were positive for the pluripotency marker Oct3/4, whereas NSCs and neurons were not, confirming the loss of pluripotency factors during differentiation. Moreover, we observed that NSCs exclusively expressed neural stem cell markers such as NeuN, Nestin, and Pax6 (Figure 2B, Figure S11, Figure S12). In contrast, neurons were positive for mature neuronal markers tubulin beta 3 and MAP2, while only a few NSCs were positive for these markers (Figure 2B, Figure S11, Figure S12). We further corroborated these findings using western blot analysis of whole cell lysates from iPSCs, NSCs, and neurons. The western blot analysis included detection of Oct3/4, MAP2, tubulin beta III, NeuN, and DCX, with lamin B1 and Ponceau red staining serving as a loading control (Table S8, Figure 2C). The western blot results were consistent with the immunostaining findings, confirming the successful differentiation of iPSCs into neurons.

Having established the neuronal differentiation system, we aimed to determine whether chromatin dynamics are affected during differentiation. To investigate this, we cultured iPSCs and NSCs on coated high-precision coverslips and labeled their chromatin using SNTT1 and dC^SiR^TP (Figure 2A). Following labeling, we replaced the buffer with growth medium and allowed the cells to recover for 1–2 hours. After this period, both iPSCs and NSCs were imaged using Airyscan microscopy. For a subset of chromatin-labeled NSCs, we replaced the medium with differentiation medium and cultured the cells for 10 days to induce differentiation into neurons before imaging.

We then identified cells with chromatin labeled by dC^SiR^TP (magenta) and performed live-cell time-lapse microscopy with Airyscan microscopy using two channels: dC^SiR^TP (magenta) and a live DNA stain (cyan). Images were acquired using Airyscan microscope (Zeiss) with 2 channels imaging and 3D z-stacks (5 stacks per channel, x-y: 48 nm and z – 170 nm) with a frame interval of Δt = 5 seconds (Figure 2D, Movie 2). Movie 2 illustrates cell movement and labeled chromatin mobility over time, along with nuclear shape changes over time. To compensate for cell movement and accurately measure chromatin mobility within the cell, we performed 3D image registration of the live-cell time-lapse movies. Representative images of labeled chromatin at the time point 1 (T1, cyan) and 100 (T100, magenta) of the time-lapse were overlaid before and after registration (Δt = 5 seconds) (Figure 2E). In the unregistered image, we observed directed chromatin motion due to cell movement, an effect eliminated by the 3D image registration (Figure 2E). This effect is further visualized by comparing chromatin tracks before and after registration (Figure 2F). We observed that after image registration, the chromatin tracks (random colors) lost their “apparent” directed motion over time, as clearly seen in Movie 3 and Figure 2F. Subsequently, we performed mean square displacement (MSD) analysis on the registered time-lapses for iPSCs (green), NSCs (blue), and neurons (magenta), and plotted the results (Figure 2G). Generally, a higher MSD curve indicates greater chromatin diffusivity. We observed (Figure 2G and 2H) that chromatin in iPSCs (14.99 x 10⁻⁵ µm²/s) is more mobile than in NSCs (8.38 x 10⁻⁵ µm²/s) and neurons (10.47 x 10⁻⁵ µm²/s). As a control for imaging system vibrations, we performed time-lapse microscopy on formaldehyde-fixed labeled cells. As shown in Movie 4, these fixed cells exhibit minimal motion or diffusion. Interestingly, NSCs and neurons showed similar diffusion rates, although chromatin in neurons appears slightly more mobile than in NSCs (Figure 2G and 2H). Previous studies have used RNA seq and ATAC seq to understand the changes in chromatin during neuronal differentiation (23) and have shown chromatin accessibility and transcription profiles are altered during differentiation. Our analysis reveals that the pluripotent state has higher chromatin mobility, and this decreases upon neural differentiation, raising the question of whether global chromatin profile changes contribute to these observed dynamics.

### Chromatin state changes during differentiation correlate with its dynamics

To analyze the DNA distribution of different cell types, we stained the DNA of iPSCs, NSCs, and neurons using DAPI. High-throughput imaging was then employed to quantify the standard deviation (Std DAPI) and total DNA amounts (SUM DAPI), which are shown in Figure 3A. Briefly, the nuclei were first segmented using Knime analytics software and the total DNA intensities (SUM DAPI) and standard deviation of DAPI (Std DAPI) measurements per cell were obtained. A higher standard deviation indicates greater variability in DNA compaction (euchromatin versus heterochromatin) across the cell population. Conversely, a lower standard deviation implies an excess of one state of chromatin. Our findings reveal that iPSCs exhibit a greater variation in standard deviation compared to NSCs and neurons, indicating a distribution of both euchromatin and heterochromatin within the iPSC population (Figure 3A). The histogram comparing total DNA amounts (SUM DAPI) across the cell types displays that iPSCs (green) include populations in the G1, S, and G2 phases, whereas NSCs contain a lower number of cells in G2 phase relative to iPSCs. Notably, neurons predominantly consist of G1 phase cells (Figure 3A). Additionally, we obtained 3D stacks of the DNA dye from all cell types and measured the volume (in µm³) and shape factor, presenting the results as boxplots in Figure 3B. During neuronal differentiation, both NSCs and neurons significantly decrease in volume compared to iPSCs and exhibit increased irregularity in shape (Figure 3B).

**Figure 3.**
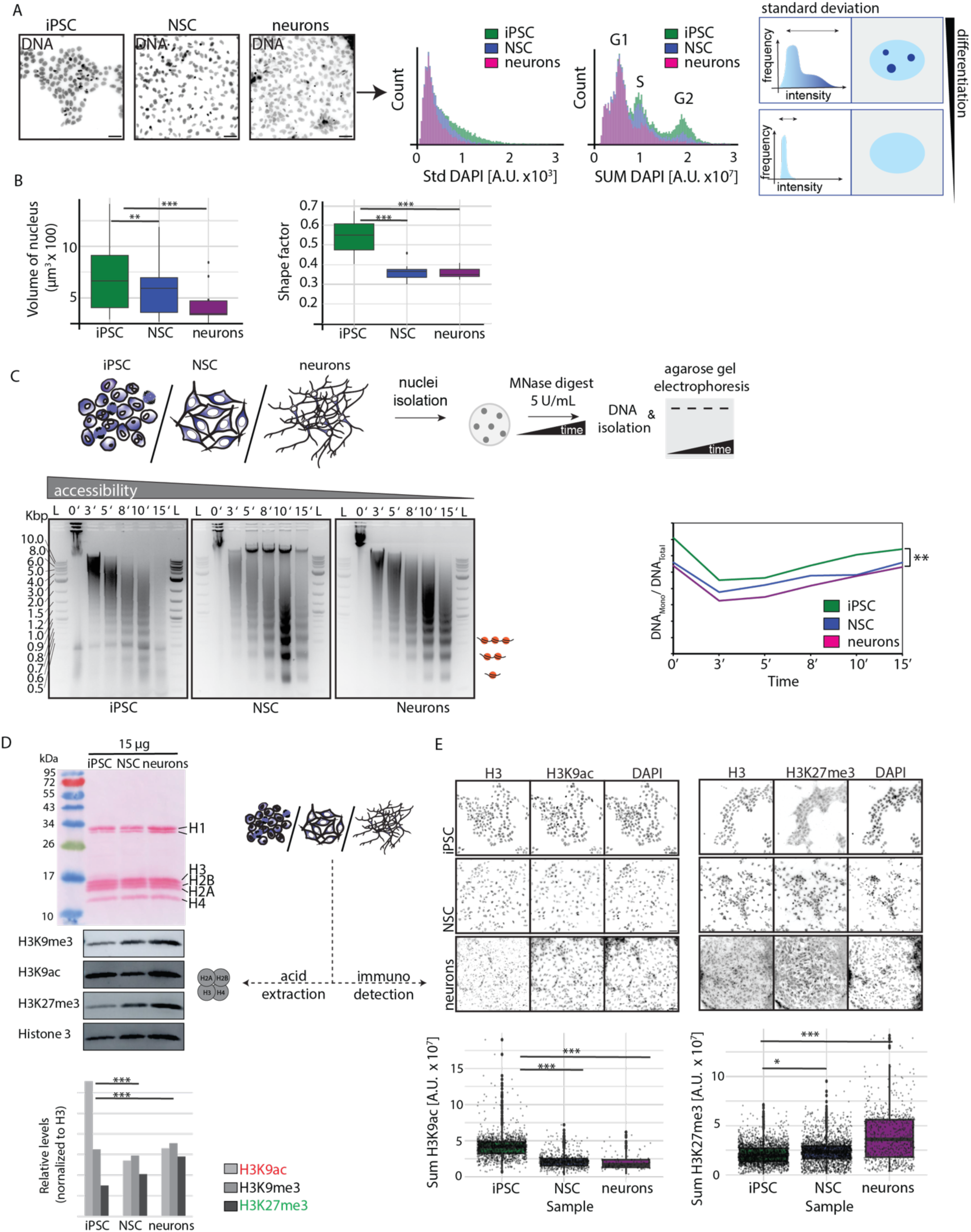
Chromatin accessibility and modifications during differentiation. **(A)** Total DNA was detected using DAPI and histograms depicting the standard deviation (std DAPI) and total DNA amount (DAPI SUM) distribution and diversity of cell populations for each cell differentiation state. Scale bars: 50 µm. **(B)** Box plots quantifying the changes in the volume of the nucleus (µm^3^) and shape factor properties of the cell differentiation states as indicated (3 biological replicates, p values: ***(p ≤ 0.001) and **(p ≤ 0.01)). **(C)** Genome-wide chromatin accessibility profiles using the MNase digestion assay (5 U/ mL) over time. The line plot shows the DNA mono/DNA total (y-axis) and time (x-axis) in minutes (2 biological replicates, p values: **(p ≤ 0.01)). **(D)** The histone fractions were isolated from different cell differentiation states using the acid extraction method and western blots were used to detect the different histone modifications. The bar plots show the relative histone modification levels normalized to the total histone H3 level (2 biological replicates, p values: ***, p ≤ 0.001). **(E)** Representative images show different histone modifications, total histone H3 and DNA (DAPI) using immunostaining. The data was analyzed and represented as a boxplot for different modifications normalized to histone H3 levels. The representative microscopy images are shown in gray scale (2 biological replicates, p values: ***(p ≤ 0.001) and *(p ≤ 0.05)). Scale bars: 50 µm.

We, then, examined whether the observed changes in global chromatin reorganization affect chromatin accessibility. To test this hypothesis, we performed a micrococcal nuclease (MNase) digestion assay to assess global chromatin accessibility in iPSCs, NSCs, and neurons. Briefly, we applied 5 U/mL MNase for various time intervals (0, 3, 5, 8, 10, and 15 minutes) to isolated genomic DNA. Subsequent agarose gel electrophoresis analysis revealed that, over time, the MNase-digested genomic DNA (detected by ethidium bromide staining) progressively transitioned from larger fragments, indicative of multiple nucleosomes, towards single nucleosome-sized fragments, observed between 10 and 15 minutes of digestion (Figure 3C). Next, we quantified the ratio of intensities of DNA fragments under 0.5 kbp (DNAmono) and total DNA (DNAtotal) within the same gel lane. The plot of the DNAmono/DNAtotal ratio (y-axis) against time (x-axis) revealed that chromatin accessibility was significantly higher in iPSCs compared to NSCs and neurons (Figure 3C). Interestingly, we found no significant differences in chromatin accessibility between NSCs and neurons (Figure 3C).

To investigate whether changes in histone modifications during neuronal differentiation affect global chromatin profiles and accessibility, we extracted histones from iPSC, NSC, and neuron cell pellets using established acid extraction protocols. A representative image of a gel analysis, visualized by Ponceau staining, revealed the presence of extracted proteins including the different histones (H1, H2A, H2B, H3, H4) (Figure 3D). We then detected by western blot analysis prominent histone modifications associated with different chromatin states using specific antibodies (Figure 3D). These included H3K9me3 (constitutive heterochromatin mark), H3K9ac (euchromatin mark), and H3K27me3 (facultative heterochromatin mark). Canonical histone H3 was also detected and its levels served as a loading control for normalization. The intensity of each histone modification band was normalized to the corresponding H3 band intensity, and the relative signal levels were plotted as bar graphs using Image lab software (Figure 3D). The western blot results indicated that global levels of H3K9ac decreased, while H3K27me3 levels increased (Figure 3D). Notably, the levels of H3K9me3 did not change significantly. To further validate these findings, we performed immunofluorescence staining for H3K9ac and H3K27me3, along with total histone H3 (Figure 3E). Representative grayscale images illustrate the localization of these histone modifications and DNA staining within the cells (Figure 3E). Quantification of the fluorescence intensity in individual nuclei was performed using Knime analytics software and presented as boxplots (Figure 3E, Table S9), corroborated the western blot results. During neuronal differentiation, H3K9ac levels decreased while H3K27me3 levels increased. iPSCs showed high H3K9ac levels, indicating active chromatin, which aligns with findings in mouse pluripotent stem cells (24). The combination of H3K4me3 and H3K27me3 creates “poised” chromatin, supporting stem cell pluripotency (25). iPSCs’ high proliferation rate leads to many cells in S phase with shorter G1 phase (26, 27). However, slower H3K27me3 restoration after replication results in diluted marks (28). As cells differentiate, iPSCs’ hyperactive chromatin state reduces, with lower H3K9ac levels. The poised chromatin becomes silenced through H3K27me3 deposition. More cells enter G1 phase (NSCs) or stop replicating (neurons), creating a repressed chromatin state with H3K27me3 marks, shown by more compact DNA (Figure 3A) and less accessible chromatin (Figure 3C) in differentiated states.

### Chromatin motion is confined during neural differentiation

To investigate whether the labeled chromatin sites exhibit different diffusion or mobility based on their chromatin state, we classified total DNA into two major categories based on pixel intensity (Figure 4A; heterochromatin - red, euchromatin - yellow) in live-cell imaging using a live-cell DNA dye. We, then, applied this classification as a mask to categorize the labeled tracks into corresponding euchromatin or heterochromatin tracks. This allowed us to track and obtain the MSD of labeled tracks located within either euchromatin or heterochromatin regions of the nucleus (Figure 4B). Consistently with previous reports (29, 30), we observed that chromatin labeled as heterochromatin is less mobile than chromatin labeled as euchromatin. As we observed in the histone modifications experiments, the decrease in the euchromatin mark (H3K9ac) during neuronal differentiation suggests that the reduced mobility of chromatin foci in heterochromatin regions contributes to the overall decrease in chromatin mobility observed in NSCs and neurons, as compared to iPSCs (see Figure 2G and 2H). However, the relative difference in mobility between heterochromatin tracks and the combined pool of all tracks was not greater in NSCs and neurons compared to iPSCs. This led us to question whether chromatin, in general, is more constrained in NSCs and neurons than in iPSCs.

**Figure 4.**
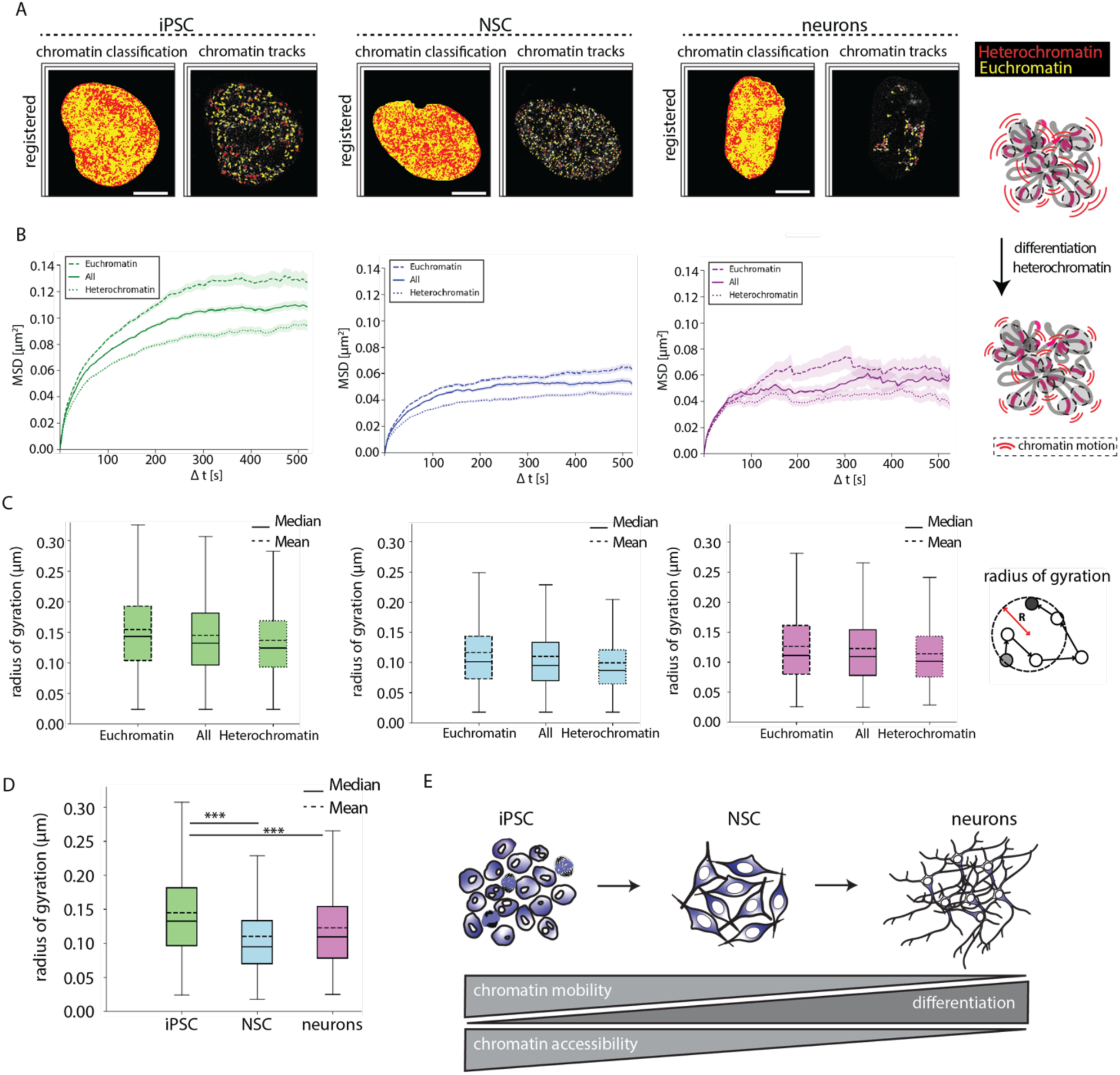
Correlation of chromatin mobility and chromatin states. **(A)** The live cell DNA dye was used to classify DNA into euchromatin and heterochromatin and the corresponding tracks of dC^SiR^TP labeled chromatin into two states - euchromatin (yellow) and heterochromatin (red). **(B)** The tracks within the chromatin states were further classified and used to perform mean square displacement analysis. The MSD (µm^2^) over time (Δt) was plotted for euchromatin, heterochromatin and combined. **(C)** Radius of gyration (µm) for euchromatin, combined (all), and heterochromatin chromatin tracks as boxplots with mean and median values for iPSCs, NSCs, neurons. **(D)** Comparison of radius of gyration (all) for iPSCs, NSCs, neurons as boxplots. **(E)** Graphical illustration of the relationship between chromatin mobility, accessibility and neuronal differentiation. Scale bars: 5 µm.

To answer this, we computed the radius of gyration (µm), which essentially quantifies how much nuclear space the chromatin can explore within a given time. We plotted the radius of gyration (µm) for euchromatin, combined (all) and heterochromatin chromatin tracks for iPSCs, NSCs, neurons and observed that overall heterochromatin is more constrained than euchromatin or both combined (Figure 4C). We then compared the radius of gyration (µm) for iPSCs, NSCs and neurons and observed that the iPSCs have larger radius of gyration (µm) than NSCs and neurons, allowing us to conclude that the chromatin in the neural differentiation state is more constrained than in the pluripotent state (Figure 4D). We propose that the restriction in chromatin accessibility during differentiation is the main cause for reduction of chromatin mobility (Figure 4E).

In conclusion, we have developed a novel STED-compatible method for labeling genomic DNA in living human induced pluripotent stem cells (iPSCs) and neural stem cells (NSCs). This method utilizes a novel silicon rhodamine-linked deoxycytidine triphosphate (dC^SiR^TP), which is delivered to live cells by the SNTT1 transporter. The dC^SiR^TP/SNTT1 combination enabled rapid chromatin labeling (within 10–15 minutes) without disrupting cell morphology, allowing for immediate high-resolution imaging and for unimpaired cell proliferation and differentiation. This approach is straightforward and highly efficient, offering significant advantages over previously reported methods, including our own (21, 22, 29). Using this system, we achieved high-resolution quantification and tracking of chromatin dynamics in living cells throughout the differentiation process from iPSCs to NSCs and ultimately to neurons. Our analyses reveal a progressive decrease in chromatin mobility that correlates with cellular differentiation state, strongly suggesting a relationship between genome dynamics and developmental potential. As NSCs are still proliferating but showed similar reduced chromatin mobility to neurons, this effect cannot be explained by cell cycle differences alone. Our observation of reduced chromatin mobility during cellular differentiation is consistent with recent studies that demonstrated using fluorescent recovery after photobleaching (FRAP) that chromatin mobility serves as a key regulatory mechanism in cellular plasticity and lineage commitment (31). The reduced chromatin mobility observed during neural differentiation likely reflects the establishment of stable transcriptional programs essential for neuronal function, providing a physical basis for cellular memory and identity maintenance. These findings provide new insights into the mechanical properties of chromatin during cell fate transitions and have implications for understanding both normal development and pathological states where chromatin organization is disrupted.

## MATERIALS AND METHODS

### Synthesis of dC^SiR^TP

dC^SiR^TP was prepared by reaction of azido-propargyloxy-triethylene glycol conjugated deoxycytidine triphosphate dC^pegN3^TP with commercially available bicyclononyne-linked silicon rhodamine fluorophore SiR-BCN in dimethylsulfoxide at room temperature. The product was isolated in 50% yield after purification by reverse-phase HPLC. Experimental conditions for the reaction, analytical data of the product (^1^H and ^13^C NMR, MS), photophysical characterization data as well as description of the biochemical experiments (in vitro DNA incorporation: PEX, PCR) are provided in supplementary information (Tables S1-S3; Figures S1-S7).

### Cells

All cells used were tested for mycoplasma and deemed free of contamination. All cell lines were authenticated by STR profiling (ATCC). All cells were grown in a humidified atmosphere of 5 % CO2 at 37°C. The details of the cell lines used are described in Table S4 To grow the human induced pluripotent stem cells clone B4 (hiPSC), surfaces were first coated with vitronectin (VTN-N) recombinant human protein (VTN) 5 ng/ml (Cat.No.: A14700, Thermo Fisher Scientific, USA) for one hour. The iPSCs clone B4 were grown in iPSC growth media containing IPS-Brew basal medium (Cat.No: 130-107-086, Miltenyi Biotec, Germany) supplemented with IPS-Brew 50x supplement (Cat.No: 130-107-087, Miltenyi Biotec, Germany) and Revita Cell™ supplement (1x) (Cat.No: A2644501, Fisher Scientific, USA) without antibiotics. For passaging of iPSCs, the cells were detached using 1x accutase (Cat.No: A6964, Sigma Aldrich Chemie, Germany) at 37 °C for three minutes, followed by centrifugation at 300 x g for three minutes. The excess media was then removed, and cells were resuspended in the growth media and seeded on vitronectin (VTN-N) coated dishes. All iPSCs cells were grown till they started forming colonies before performing experiments (Figure S8). The iPSCs were differentiated in vitro to neural stem cells (NSCs) (Figure S9) and the detailed protocol and media compositions are described in supplementary information section 6 and Table S7.

### Immunofluorescence

Cells on coverslips were fixed with 3.7% formaldehyde for 10 minutes. Cells were permeabilized with 1x PBS supplemented with 0.7% Triton X-100 for 20 minutes. After washing three times with PBS-T for five minutes, cells were blocked with 1% BSA/PBS for 30 minutes and subsequently incubated for an hour with the primary antibody, followed by three washing steps with PBS-T. Cells were incubated with a secondary antibody for 45 minutes followed by three washing steps with PBS-T. DNA was counterstained with 4′,6-diamidino-2-phenylindole (DAPI) (Cat. No. 6335.1, Carl Roth, Germany) for 10 minutes. Coverslip was mounted onto glass slides using Vectashield before proceeding to imaging.

iPSCs were seeded and cultured on glass coverslips until the colonies were obtained. The media was replaced with media containing 10 µM EdU and pulsed for 10 minutes (Table S5). Subsequently, the cells were fixed with 3.7% formaldehyde for 10 minutes. The samples were used to detect pluripotent stem cell markers (Sox2, Oct3/4) and EdU was detected using Click-IT chemistry (Table S8, Figure S8).

### Micrococcal nuclease chromatin accessibility assay

A total of 10×10^6^ cells were harvested, and cell pellets were resuspended in a hypotonic lysis buffer (10 mM Tris, pH 7.5, 10 mM NaCl, 3 mM MgCl2 and 0.5% NP-40) for five minutes on ice. Cell nuclei were pelleted (five minutes, 120 × g), washed once in digestion buffer (50 mM Tris pH 8, 5 mM CaCl2) and resuspended in digestion buffer. Microccocal nuclease (MNase) (cat# EN0181, Thermo Fisher Scientific, Waltham, MA, USA) was diluted in MNase storage buffer (20 mM TrispH7.6, 50 mM NaCl, 50% glycerol), 5 U/ mL, were added to the nuclei in a final volume of 100 µl. Reactions were incubated for different timepoints at 37 °C, stopped by the addition of stopping buffer (20mM EDTA, 0.4% SDS, 0.5 mg/ml proteinase K) and incubated at 50 °C overnight. DNA was recovered by adding 0.1 volumes of 3 M NaCl and 2 volumes of 100% ethanol and storage at –20 °C for an hour. After centrifugation (15 minutes, 15 000 × g) and washing with 70% ethanol, the DNA pellet was air-dried and resuspended in 50 µl ddH2O. Equal amounts of DNA were loaded on a 1.5% agarose gel and electrophoresed for 45 min at 100 V. Gels were stained with ethidium bromide in 1× TAE buffer (Tris-acetate-EDTA) and imaged using an AI600 imager. Integrated intensities of monomeric DNA and total DNA in each lane were measured with ImageJ (Table S9) and plotted as a DNA monomer/DNA total ratio.

### Western blot

The cells were cultured as described above. The cells were detached using 2 mL of accutase (Cat.No: A6964, Sigma Aldrich Chemie, Germany) and counted and 2 x 10^6^ cells were collected into a tube. The lysis buffer NP-40 was supplemented with the protease inhibitors: 1mM PMSF (Phenylmethylsulfonyl fluoride, Cat.No: 6367.2, Carl-Roth, Germany), 1 mM AEBSF (4-(2-Aminoethyl) benzyl sulfonyl fluoride hydrochloride, Cat.No: A1421.0100, VWR, USA), 1 mM E64 (Cat.No: E3132, Sigma-Aldrich, USA) and 1 nM Pepstatin A (Cat.No: 77170, Sigma-Aldrich, USA). Cells were homogenized in cell lysis buffer containing 150 mM NaCl (Cat.No.: 0601.2, Carl Roth, Karlsruhe, Germany), 200 mM TrisCl pH 8 (Cat.No.: A1086.500, Diagonal, Münster, Germany), 5 mM EDTA (Cat.No.: 8040.2, Carl Roth, Karlsruhe, Germany) 0.5% NP-40 (Cat.No.: 74385, Sigma-Aldrich Chemie GmbH, Steinheim, Germany and incubated for 30 minutes at 4°C. The lysates were cleared by centrifuging for 30 minutes at 14,000 g at 4°C. Protein concentration was determined using Pierce™ 660 nm Protein Assay Reagent (cat# 22660, Thermo Fisher Scientific, Waltham, MA, USA) according to the manufacturer’s instructions. Protein lysates were denatured in 6x protein loading dye (560 mM Tris-HCl pH 6.8, 60 mM DTT, 6 mM EDTA, 30% glycerol and 0.6% bromophenol blue) for five minutes at 95 °C. Denatured protein lysates were separated by 10% SDS-PAGE and transferred onto 0.45 μm nitrocellulose membranes (GE Healthcare/Whatman; Catalog #10600002). The membranes were blocked with 3% low-fat milk for 30 minutes and subsequently incubated with primary antibodies diluted in blocking solution for 2 hours at 4 °C (Table S8). After washing three times with PBS-T for five minutes each, membranes were incubated with secondary antibodies for 1 hour at room temperature (Table S8). Following two wash steps with PBS-T, protein bands on the membranes were visualized using an AI600 imager (GE Healthcare) (Table S6).

### Acid extraction of histones

The histone proteins were acid-extracted using the protocol described in (32). Around 10^6^ cells were taken from each cell type and centrifuged. The cells were washed with 1x PBS before resuspending in 1 ml of hypotonic lysis buffer (10 mM Tris-Cl pH 8.0, 1 mM KCl, 1.5 mM MgCl2, and 1 mM DTT. Protease inhibitors were supplemented before use). The cells were incubated for 30 minutes on a rotator for hypotonic swelling and lysed by mechanical shearing. The nuclei were pelleted using centrifugation at 10,000 x g for 10 minutes at 4 °C. The nuclei were resuspended in 0.4 N H2SO4 and incubated overnight. The cells were then carefully resuspended using 200 µl pipette tips. The nuclei debris was removed by centrifuging the mixture at 16000 x g at 10 minutes, and the supernatant was transferred to a new tube. Around 132 µl of 100 % TCA (Trichloroacetic acid) was added dropwise to make a final concentration of 33% and inverted several times to precipitate the histones. The samples were incubated on ice for 30 minutes. The lysate was centrifuged at 16000 x g for 10 minutes at 4 °C. The supernatant was removed, and the pellet was washed twice with ice-cold acetone to remove the acid. After drying the histones at room temperature for 20 minutes, they were dissolved in 50 µl of ddH2O. The concentration of the proteins was measured using Pierce™ 660nm Protein Assay Reagent (cat# 22660, Thermo Fisher Scientific, Waltham, MA, USA). Protein lysates were denatured in 6x protein loading dye (560 mM Tris-HCl pH 6.8, 60 mM DTT, 6 mM EDTA, 30% glycerol and 0.6% bromophenol blue) for five minutes at 95 °C. 10 µg of denatured protein lysates were resolved using 10% SDS-PAGE and transferred onto 0.45 μm nitrocellulose membranes (GE Healthcare/Whatman; Catalog #10600002). The membranes were blocked with 3% low-fat milk for 30 minutes and subsequently incubated with primary antibodies diluted in blocking solution for 2 hours at 4°C (Table S8). After washing three times with PBS-T for five minutes each, membranes were incubated with secondary antibodies for 1 hour at room temperature (Table S8). Following two wash steps with PBS-T, protein bands on the membranes were visualized using an AI600 imager (GE Healthcare) (Table S6).

### Chromatin labeling using SNTT1

The iPSCs and NSCs were cultured as described above. The cells were split onto glass bottom culture dishes for chromatin labeling, and incubated with tricine buffer (pH 7.4) containing 10 µm SNTT1 (Cat.No: SCT064, Merck Millipore, Germany) + 10 µm dC^SiR^TP for 15 minutes at 37 °C (Table S5). The cells were then washed with 1 x PBS to remove excess nucleotides. The cells were subsequently incubated in media containing 10 µm of Abberior DNA live 590 dye (Cat.No: LV590-0143-50UG, Abberior, Germany) or Abberior tubulin live 610 dye (Cat.No: LV610-0141-50UG, Abberior, Germany) for three hours (Table S5).

### Super resolution time lapse microscopy

The chromatin and total DNA was labeled in live cells as described above. Following the labeling, the cells on the glass bottom were used to perform live/fixed cell time lapse microscopy for chromatin mobility measurements. Briefly, time lapse imaging was performed within 1-2 hours after labeling of the cells using a Zeiss LSM 900 Airyscan system (Zeiss, Germany) equipped with a C Plan-Apochromat 63x oil objective and Axiocam 820 mono SONY IMX541 CMOS camera with Airyscan detector for high-speed stack acquisition. The details of the microscope are described in Table S6. The DNA dye was used to find the focal plane of cells to minimize bleaching in the chromatin (dC^SiR^TP) channel. The DNA dye and dC^SiR^TP were excited using 561 nm and 640 nm lasers respectively. All chromatin motion timelapse microscopy was performed using two channel imaging and 5 z-stacks per channel with a lateral (x-y) resolution of 48 nm and axial resolution (z) of 170 nm with 5 s interval. The raw data was processed to obtain super resolution time lapse movies using Airyscan joint convolution with 20 iterations of processing (Zeiss) (Table S9). The time lapse imaging was performed at 37 °C with a humidified atmosphere using an environmental chamber (Table S6). The standard protocol for examining chromatin mobility in dC^SiR^TP labeled nuclei proceeded in the following manner: first, a reference overview image of dC^SiR^TP and Abberior DNA live 590 dye using wide-field mode was collected from a single focal plane corresponding to the middle of the nucleus. Second, while maintaining the same central focal plane, a time series (frame interval of 5 s, 5 z-stacks, 200 frames) and 3D volume with fewer stacks were captured to minimize photo toxicity. The imaging was performed over multiple experiments to have reproducibility and sufficient replicates.

Fixed cell imaging was performed as an imaging control, where the cells were labeled with chromatin (dC^SiR^TP) and Abberior DNA live 590 dye. Post labeling, the cells were fixed using 3.7 % formaldehyde for 10 minutes at room temperature. The imaging was performed as described above.

For fixed cell STED super resolution imaging, the iPSCs were cultured as described above and seeded on high precision (18 x 18 mm) square coverslips. The cells were labeled with chromatin (dC^SiR^TP) and Abberior DNA live 590 dye. The cells were allowed to recover from the labeling for 2-3 hours and fixed using 3.7 % formaldehyde for 10 minutes at room temperature. The coverslip was then mounted using Vectashield before proceeding to imaging using STED 775 QUAD Scan microscope (Abberior Instruments) (Table S6).

### 3D motion analysis of chromatin

To decouple the movement of the chromatin structures from the movement and deformation of the cell, joint affine and non-rigid 3D image registration was performed using the method in (33) adapted for 3D registration. To ensure that image registration does not affect the motion of the chromatin structures, registration was performed using the DNA (DAPI) channel images. The transformation parameters obtained were subsequently applied to the chromatin (labeled nucleotide) channel. Each frame of a live-cell image sequence was registered to the first frame.

The 3D chromatin structures within cell nuclei were tracked in the registered 3D live-cell fluorescence microscopy image sequences using a probabilistic particle tracking method (34). The method combines Kalman filtering with particle filtering and uses sequential multi-sensor data fusion and separate sensor models to integrate multiple measurements. Detection-based and prediction-based measurements were obtained by elliptical sampling (35). For detecting 3D chromatin structures, the spot-enhancing filter (SEF) (36) was used, which uses the Laplacian of Gaussian (LoG) filter followed by thresholding and computing local maxima.

To quantify the mobility of the 3D chromatin structures, a mean-square displacement (MSD) analysis was performed (37). For this, the MSD over time intervals Δt was calculated for each trajectory and then averaged. Both the diffusion model and the anomalous diffusion model were fitted to the averaged MSD values yielding the diffusion coefficient D [μm^2^s^-1^] and the anomalous diffusion coefficient α (motion type parameter). To improve the robustness of the motility analysis, only trajectories with a minimum time duration of about 50 seconds (10-time steps) were considered. To further characterize the 3D motion of the chromatin structures, we computed the radius of gyration ((38)) for each trajectory.

In addition, we analyzed the mobility of the 3D chromatin structures with respect to the 3D nuclear landscape. We divided the 3D cell nucleus into two compaction classes based on the intensity values of the DAPI channel using the method in (39). For each frame of an image sequence, first the cell nucleus was segmented and then intensity-based classification using a Gaussian Mixture Model was performed. Computed trajectories were assigned to the compaction class containing the majority of the track points.

## ACKNOWLEDGEMENTS

We thank James Adjaye for providing us with the human induced pluripotent stem cells used in this study. We also would like to thank Bodo Laube for providing antibodies and valuable insights. STED images were acquired at the Center for Advanced Light Microscopy (CALM) at the LMU Munich.

## DATA AVAILABILITY STATEMENT

All data are available at the TU Data lib at (https://tudatalib.ulb.tu-darmstadt.de/handle/tudatalib/4441) Renewable biological materials and software will be made available upon request from M. Cristina Cardoso, Michal Hocek and Karl Rohr. All experimental data including mass spectrometry and NMR spectra are presented in supplementary files.

## 1. Synthesis

### 1.1. General remarks

Solvents and reagents were purchased from commercial suppliers (Sigma-Aldrich, AlfaAesar, Spirochrome). The reactions were monitored by thin-layer chromatography (TLC) using Merck silica gel 60 F254 plates and visualized by UV (254 nm). Purification by column chromatography was performed using silica gel (40–63 µm). Separations of dC^SiR^TP were performed using HPLC (Waters modular HPLC system) on a column packed with 10 μm C18 reversed phase (Phenomenex, Luna C18). NMR spectra were measured on Bruker AVANCE 500 (^1^H at 500.0 MHz, ^13^C at 125.7 MHz, ^31^P at 202.3 MHz,) NMR spectrometer in D2O at 25 °C. Chemical shifts (in ppm, δ scale) were referenced to *t*BuOH = 1.24 ppm. Coupling constants (*J*) are given in Hz, chemical shifts in ppm (δ scale). Complete assignment of all NMR signals was achieved by using a combination of H,H-COSY, H,C-HSQC and H,C-HMBC experiments. Mass spectra and high-resolution mass spectra were measured by ESI ionization technique and spectra were measured on a LTQ Orbitrap XL spectrometer (ThermoFisher Scientific). dC^pegN3^TP was prepared according to a published procedure (1).

### 1.2. Silicon rhodamine modified deoxycytidine-5’-*O*-triphosphate (dC^SiR^TP)

A solution of SiR-BCN (2.5 mg, 3.23 µmol, in 420 µl DMSO) was added to a solution of dC^pegN3^TP (3.5 mg, 3.23 µmol, in 140 µl HPLC grade H2O) and the reaction mixture was stirred overnight at room temperature. Then, the DMSO was removed by lyophilization, and the reaction mixture was purified by reverse-phase HPLC (eluent: H2O/0.1 M triethylammonium acetate/acetonitrile 37/20/43). Collected fractions were lyophilized, and the excess of the buffer was removed by repetitive freeze-drying from water. The product was obtained as a pale yellow solid (2.84 mg, 50%).

^1^H NMR (500.0 MHz, D2O, ref(*t*BuOH) = 1.24 ppm): 0.46, 0.58 (2 × s, 2 × 3H, CH3Si); 0.66 – 0.78 (bm, 2H, H-5a,6a-CPCOT); 0.86 (bm, 1H, H-6-CPCOT); 1.10 – 1.24 (bm, 2H, H-5b,7b-CPCOT); 1.27 (t, 27H, *J*vic = 7.3, CH3CH2N); 1.91, 2.02 (2 × bm, 2 × 1H, H-5a,7a-CPCOT); 2.18, 2.38 (2 × bm, 2 × 1H, H-2ʹ); 2.54 –2.66 (bm, 1H, H-4b,8b-CPCOT); 2.69 – 2.83 (bm, 1H, H-4a,8a-CPCOT); 2.95 – 3.15 (bm, 2H, NCH2CH2O); 3.15 – 3.30 (bm, 30H, (CH3)2N, CH3CH2N); 3.40 – 3.56, 3.56 – 3.71, 3.71 – 3.77, 3.77 – 3.85, 3.92 (5 × bm, 22H, OCH2CH2O, NCH2CH2O, triazole NCH2CH2O, CH2O); 4.09 – 4.23 (bm, 3H, H-4ʹ,5ʹ); 4.30 – 4.41 (bm, 4H, CH2C≡C, triazole NCH2CH2O); 4.55 (bm, 1H, H-3ʹ); 6.12 (bm, 1H, H-1ʹ); 6.49, 6.60 (2 × bm, 2 × 1H, H-2,8-DHBS); 6.90 – 7.04 (bm, 2H, H-1,9-DHBS); 7.20 – 7.29 (bm, 2H, H-4,6-DHBS); 7.69 (bm, 1H, H-3-Ph); 7.95 – 8.03 (H-5,6-Ph); 8.05 (bm, 1H, H-6).

^13^C NMR (125.7 MHz, D2O, ref(*t*BuOH) = 30.29 ppm): -2.79, -0.31, -0.28 (CH3Si); 8.92 (CH3CH2N); 17.65, 17.69 (CH-6-CPCOT); 19.28, 19.36, 19.78, 19.81 (CH-5a,6a-CPCOT); 21.79, 22.23 (CH2-5,7-CPCOT); 23.04, 25.87 (CH2-4,8-CPCOT); 40.14 (CH2-2ʹ); 40.50 (NCH2CH2O); 40.70, 40.75, 40.79 ((CH3)2N); 40.84 (NCH2CH2O); 47.36 (CH3CH2N); 48.46 (triazole-NCH2CH2O); 59.19 (CH2C≡C); 63.80 (CH2O); 65.90 (CH2-5ʹ); 69.45, 69.65, 69.79, 70.07, 70.22, 70.78 (OCH2CH2O, triazole-NCH2CH2O, NCH2CH2O); 71.21 (CH-3ʹ); 77.61 (CH2C≡C); 86.27 (d, *J*C,P = 7.3, CH-4ʹ); 87.10 (CH-1ʹ); 92.40 (C-5, CH2C≡C); 114.28, 114.42, 114.49 (CH-2,8-DHBS); 121.02, 121.07, 121.16 (CH-4,6-DHBS); 128.12 (CH-5-Ph or CH-6-Ph); 128.58 (C-9a,10a-DHBS); 129.28 (CH-3-Ph); 129.78 (CH-5-Ph or CH-6-Ph); 135.64 (C-4-Ph); 140.21, 140.25, 140.53 (CH-1,9-DHBS); 140.63 (C-1-Ph); 145.69 (CH-6); 147.64, 147.67, 147.79 (C-4a,5a-DHBS); 153.85, 153.95 (C-3,7-DHBS); 155.95 (C-2); 158.69 (OCONH); 165.20 (C-4); 169.88 (CONH-4-Ph); 173.66 (COO-1-Ph); C-2-Ph, C-10-DHBS and C-3a,8a-CPCOT not detected.

^31^P{^1^H} NMR (202.4 MHz, D2O): -22.52, -10.90, -10.18 (3 × bm).

HRMS (ESI^-^): calculated for C62H82O23N10P3Si: 1455.4542; found 1455.4529.

## 2. Photophysical properties

### Molar absorption coefficients

Absorption coefficients were measured using 1 mL quartz cuvettes.

The absorption coefficients were calculated using the following equation

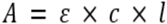

where *A* is the absorbance of the sample, *ε* is the absorption coefficient, *c* is the exact concentration of the sample and *l* is the length of the path that the light travels through the cuvette.

### Fluorescence quantum yields

Relative determination of the fluorescence quantum yields (Ф) was performed using Cresyl violet perchlorate in MeOH (Ф = 0.54) as a reference. The absorbance of sample solutions was kept below 0.10 to avoid inner filter effects. The quantum yields were calculated using the following equation (2).

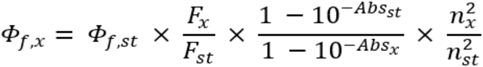

where Φf is the quantum yield, F is the integrated fluorescence intensity, Abs is the absorbance of the solution at the excitation wavelength, n is the refractive index of the solvent, and the subscripts x and st stand for the sample and standard respectively. Samples were excited at 600 nm and the emission spectra were recorded in the range of 620-800 nm.

## 3. Biochemistry

### 3.1. Enzymatic synthesis of modified DNA bearing one silicon rhodamine modification by primer extension (gel analysis)

The reaction mixture (20 μl) contained FAM labeled primer Prim^PEX^-FAM (for sequence see Table S2; 3 μM, 1 μL), template Temp^PEX^ (for sequence see Table S2; 3 μM, 1.5 μL), KOD XL DNA polymerase (0.25 U/µl, 0.3 μL), natural dGTP (4 mM, 0.6 μL), either natural dCTP (4 mM, 0.3 μL) or dC^SiR^TP (4 mM, 0.3 μL) in corresponding reaction buffer (10x, 2 μL) supplied by the manufacturer. The reaction mixture was incubated for 30 min at 60°C in the thermal cycler. The reaction was stopped by the addition of PAGE stop solution (20 µl) and the reaction mixture was denatured at 95°C for 5 min and analyzed using 12.5% denaturing PAGE. The PAGE gel was visualized by a fluorescent scanner (Figure S2A).

### 3.2 Preparation of modified dsDNA bearing one silicon rhodamine modification by primer extension (semi-preparative scale)

The reaction mixture (50 µl) containing primer (for sequence see Table S2; Prim^PEX^, 100 µM, 2.5 µl), template (for sequence see Table S2; Temp^PEX^, 100 µM, 2.5 µl), dGTP (4 mM, 1 µl), dC^SiR^TP (4 mM, 2.5 µl), KOD XL DNA polymerase (Recombinant *Thermococcus kodakaraensis*, Cat.No: 71087, Merck Millipore, Germany) (2.5 U/µl, 1.5 µl) in corresponding reaction buffer (10×, 5 µl) supplied by the manufacturer. The reaction mixture was incubated for 75 min at 60°C in a thermal cycler. The reaction was stopped by cooling at 4°C. The modified dsDNA was purified using spin columns (QIAquick® Nucleotide Removal Kit, QIAGEN) and eluted by milli-Q water. Prepared dsDNA (Table S3) was used for the measurement of photophysical properties (Table S1, Figure S1). The product was analyzed using MALDI-TOF MS; calculated for [M]: 6942.2 Da; found: 6940.9 Da.

### 3.3. Enzymatic incorporation of dC^SiR^TP by polymerase chain reaction

The reaction mixture (20 µl) contained primer (for sequences see Table S2; Prim1^PCR^ and Prim2^PCR^, 10 µM, 1 µl of each), template (Temp^PCR^, 1 nM, 0.5 µl), natural dNTPs (dATP, dGTP, dTTP, 0.4 mM each, 1.5 µl) and either dCTP (0.4 mM, 1.5 µl), dC^SiR^TP (0.4 mM, 1.5 µl) or mixture of dC^SiR^TP with natural dCTP (0-95% of dCTP), KOD XL DNA polymerase (2.5 U/µl, 0.5 µl) and corresponding reaction buffer (10×, 2 µl) supplied by the manufacturer. After the initial denaturation for 3 min at 94 °C, 40 PCR cycles were run under the following conditions: denaturation for 20 sec. at 94°C, annealing for 30 sec. at 58 °C, extension for 30 sec. at 72 °C. The reaction was stopped by cooling to 4 °C. The PCR products were analyzed by agarose gel electrophoresis in 2% agarose gel stained with GelRed™ (Biotium, agarose gels are shown in Figure S2B).

## 4. Copy of ^1^H, ^13^C and ^31^P NMR spectra (dC^SiR^TP)

## 5. Copy of mass spectra

## 6. Generation of neural stem cells (NSC) and neural differentiation

hiPSC A4 and B4 were seeded at low density on p60 (Cat.No: 83.3901, Sarstedt, Germany) in hiPSC growth media (Methods: Cells) coated with vitronectin VTN 10 ng/ml (Cat.No.: A14700, Thermo Fisher Scientific, USA) for one hour. The cells were then allowed to grow for 6-7 days to form colonies. The cells were then treated with 1 mg/ml collagenase type IV (Cat.No: C4-28, Sigma Aldrich Chemie, Germany) in DMEM for 30 minutes until the colonies were detached and floating. The collagenase IV was deactivated using hiPSC growth media. The colonies with the media were then transferred to a 15 ml conical tube using a wide bore tip and were allowed to settle down for 10-15 minutes. The supernatant was aspirated slowly and resuspended in embryoid bodies formation media (Table S7) and were grown for 10 days in the media. The embryoid bodies were then transferred for attachment onto a 6 well plate coated with poly-L-ornithine / laminin in ITSF media and cultured for 10 days (Table S7). The attached embryoid bodies were disrupted using a glass hook and the cell clumps were transferred into a culture dish with no coating and resuspended into neurosphere media (Table S7). The neurospheres were cultured for another seven days. The neurospheres were then transferred into a 15 ml conical tube and were allowed to settle down for 10 - 15 minutes. The supernatant was then removed and 1 mL of accutase was added to the tube to dissociate the neurospheres. The cells were then resuspended in 5 mL neurosphere media, followed by centrifugation at 300 r.c.f for 5 minutes. The supernatant was removed, and cells were resuspended in 5 mL NSC media and transferred to poly-L-ornithine / laminin coated dishes. The NSCs were cultured and split every 5 days until the NSCs were obtained.

For differentiation experiments, NSCs were seeded on poly-L-ornithine / laminin coated dishes at 10^5^ cells per cm^2^ density. Next day, the media was replaced with neuronal differentiation media (Table S7). The cells were allowed to differentiate for 10 days before proceeding with live cell imaging.

## 7. Supplementary movies

**Movie 1**. Live cell time lapse imaging of chromatin labeling in iPSC. Scale bar: 50 µm.

**Movie 2.** Live cell time lapse imaging of labeled chromatin in iPSC, NSC, neurons. Scale bar: 5 µm.

**Movie 3.** Registered and non-registered timelapses of labeled chromatin in iPSC, NSC, neurons. Scale bar: 5 µm.

**Movie 4.** Live and fixed cell time lapse imaging in iPSC. Scale bar: 50 µm

**Figure S1.**
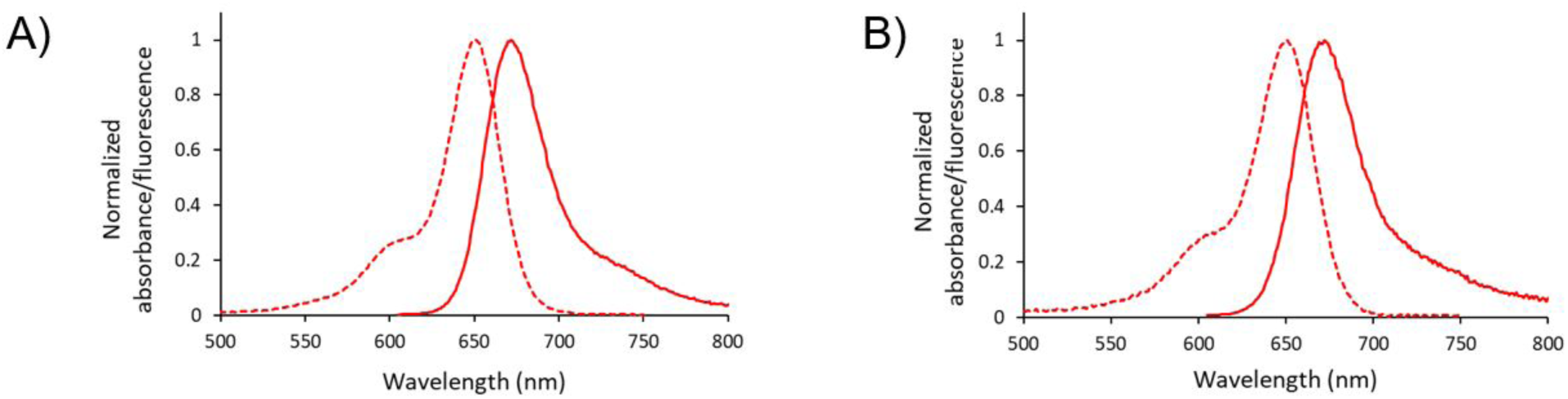
Absorption and fluorescence spectrum of **dC^SiR^TP** (A) and **DNA19_1C^SiR^** (B) in PBS.

**Figure S2.**
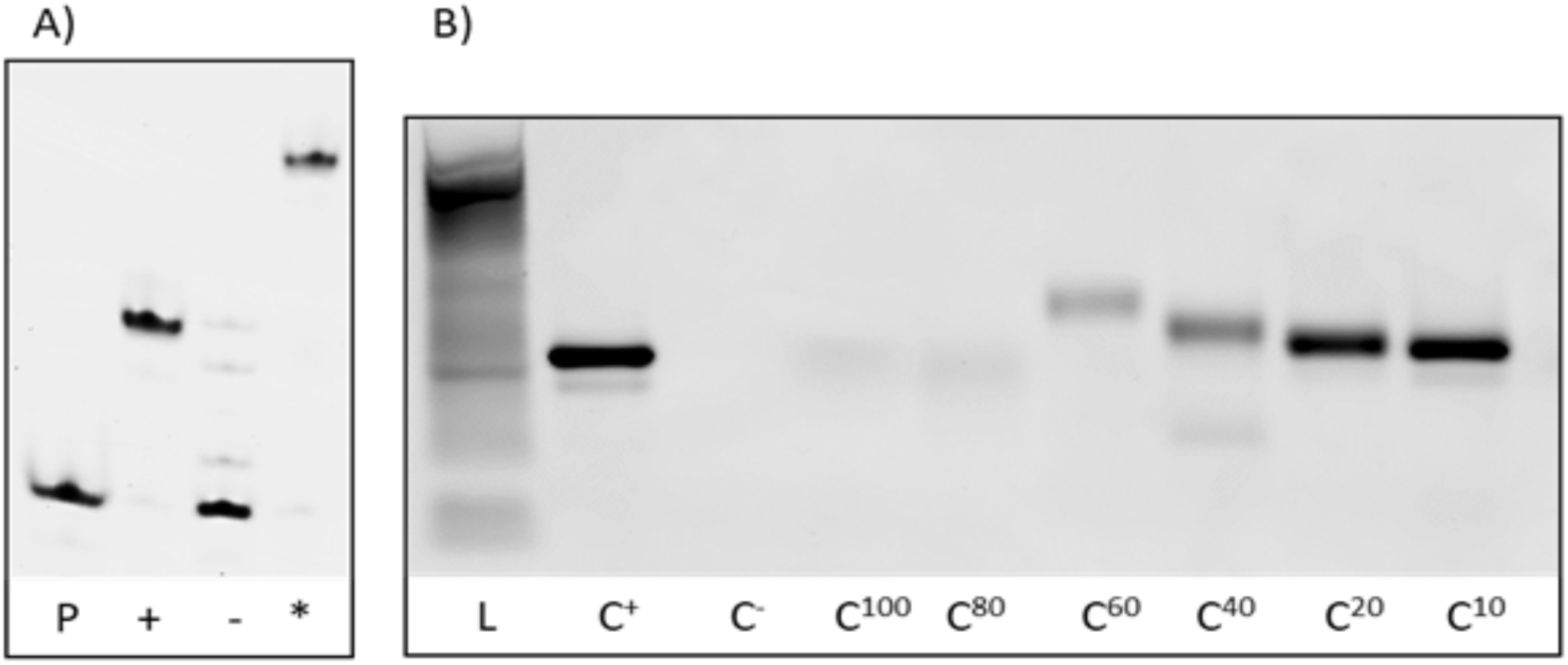
**A)** Page analysis of PEX using KOD XL polymerase, dC^SiR^TP and template Temp^PEX^. Primer (P), positive control (+, PEX with all natural dNTPs), negative control (-, PEX in absence of dCTP), PEX with dC^SiR^TP (*). **B)** Agarose gel electrophoresis analysis of PCR amplification of 98 bp template Temp^PCR^ with KOD XL DNA polymerase using dC^SiR^TP. DNA ladder (L), primer (P), positive control (C^+^, PCR with natural dNTPs), negative control (C^-^, PCR in absence of dCTP), PCR with a mixture of modified nucleotide with natural dCTP (C^100^-C^10^, the content of modified nucleotide decreases from 100% to 10%).

**Figure S3.**
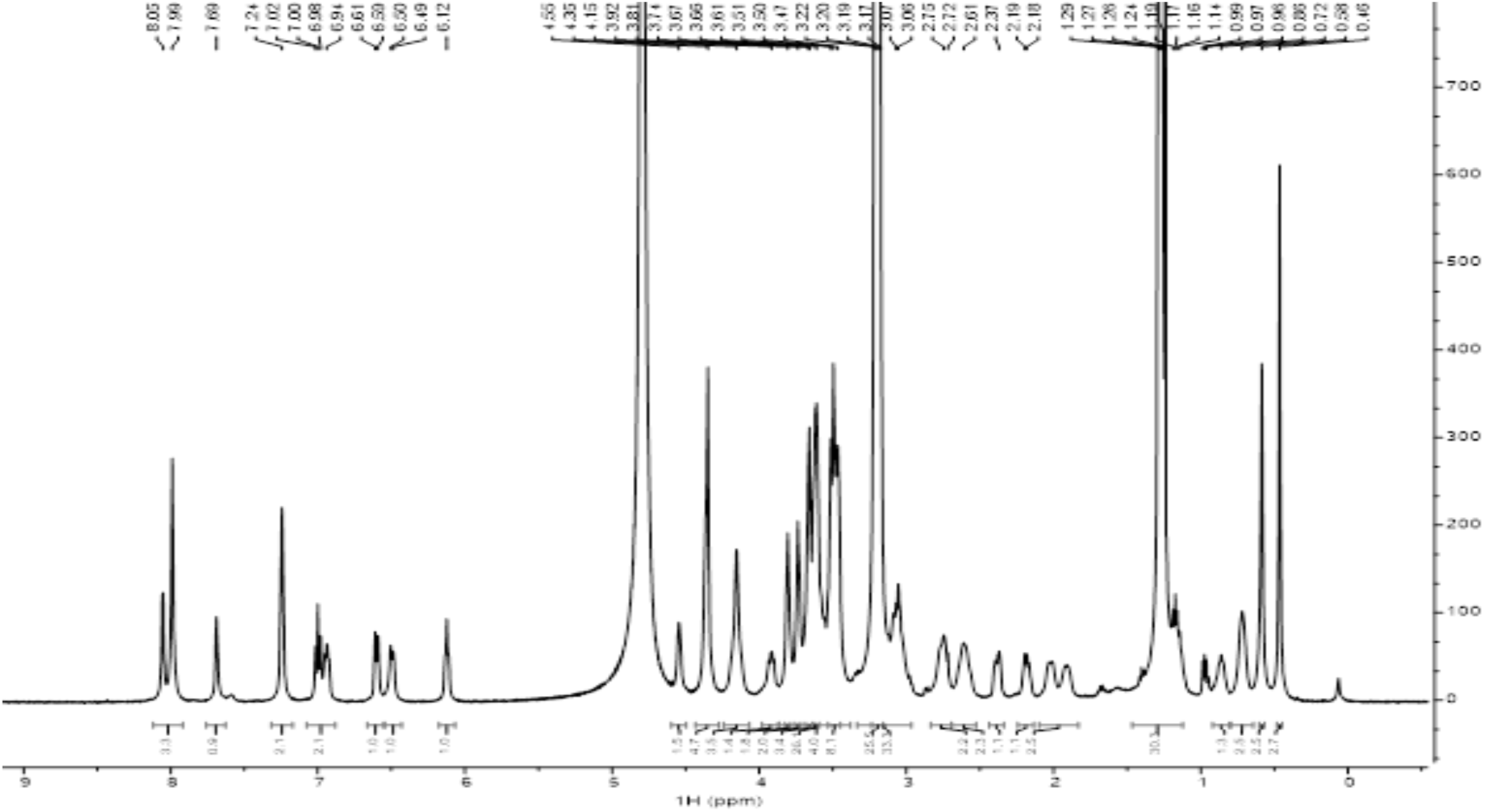
^1^H NMR spectrum of dC^SiR^TP in D2O.

**Figure S4.**
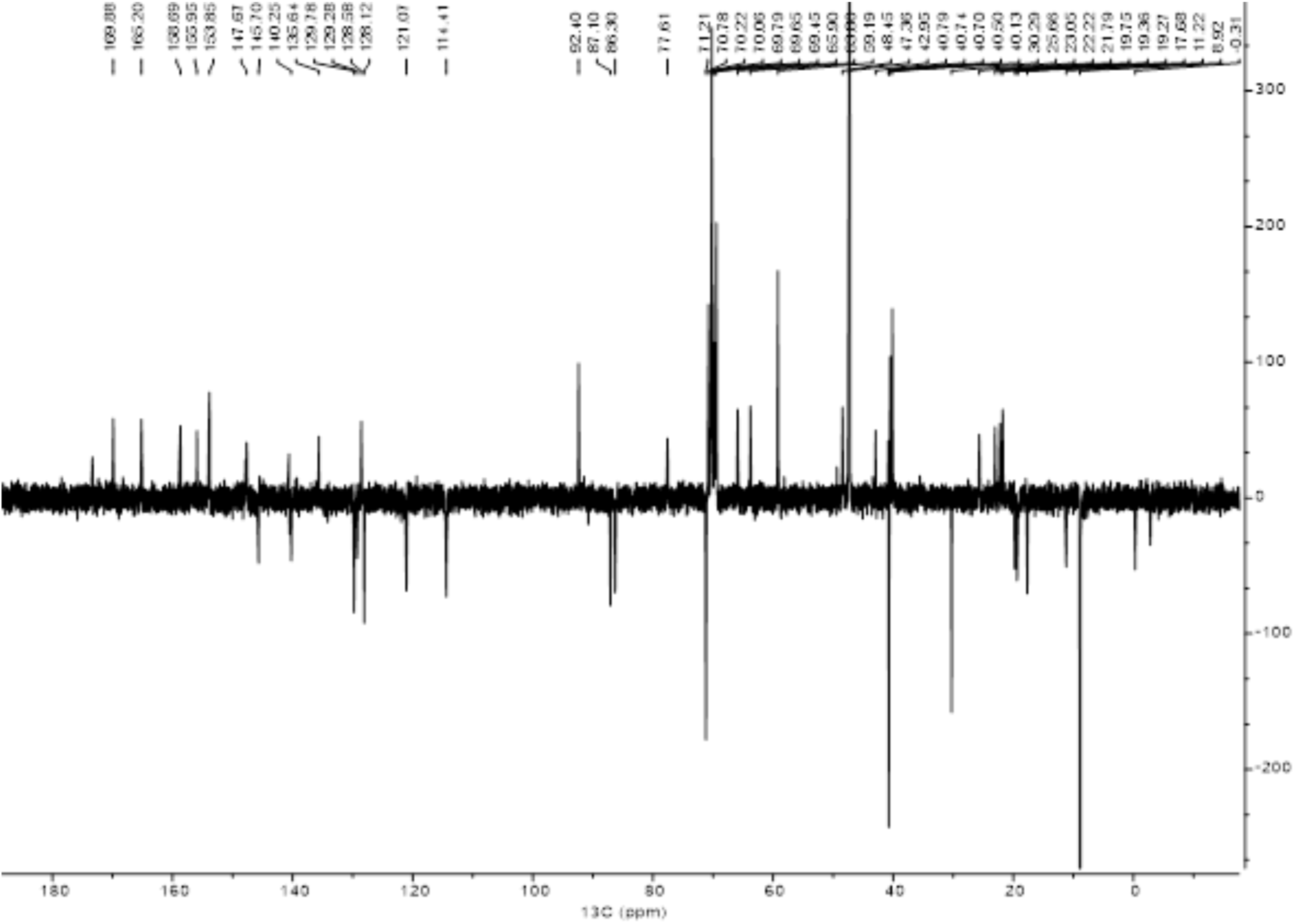
^13^C NMR spectrum of dC^SiR^TP in D2O.

**Figure S5.**
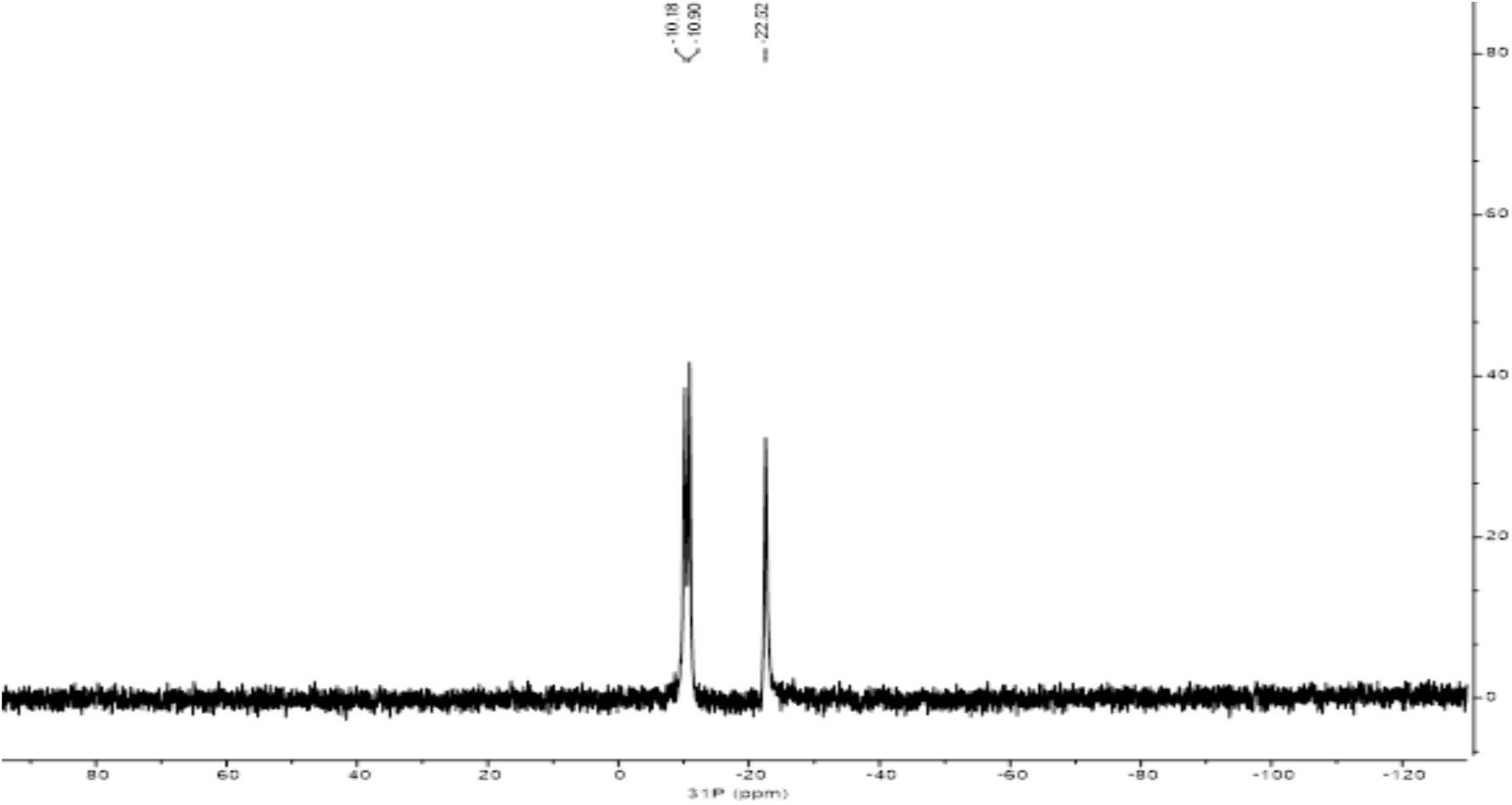
^31^P{^1^H}NMR spectrum of dC^SiR^TP in D2O.

**Figure S6.**
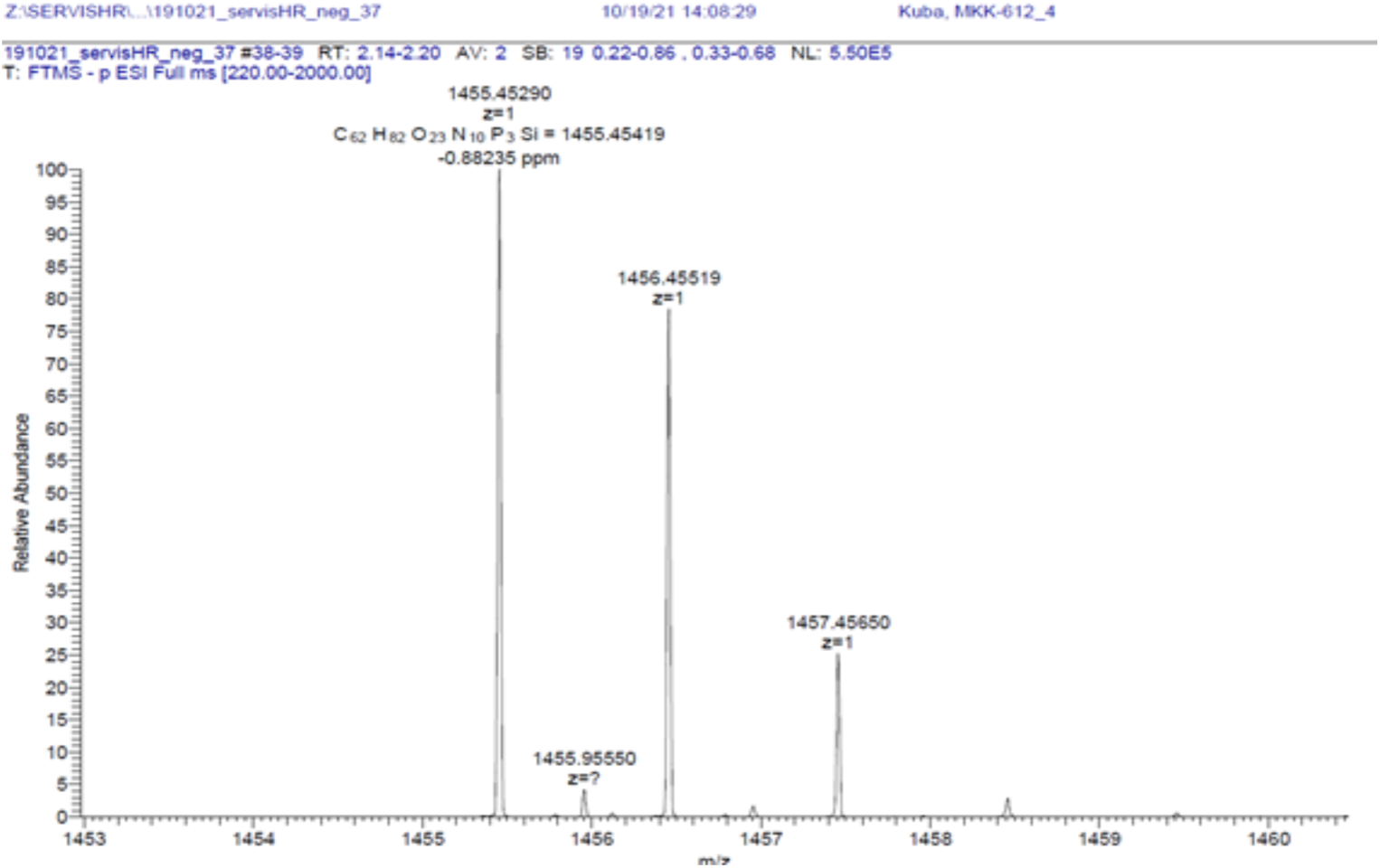
ESI MS (neg. mode) high resolution spectra of dC^SiR^TP.

**Figure S7.**
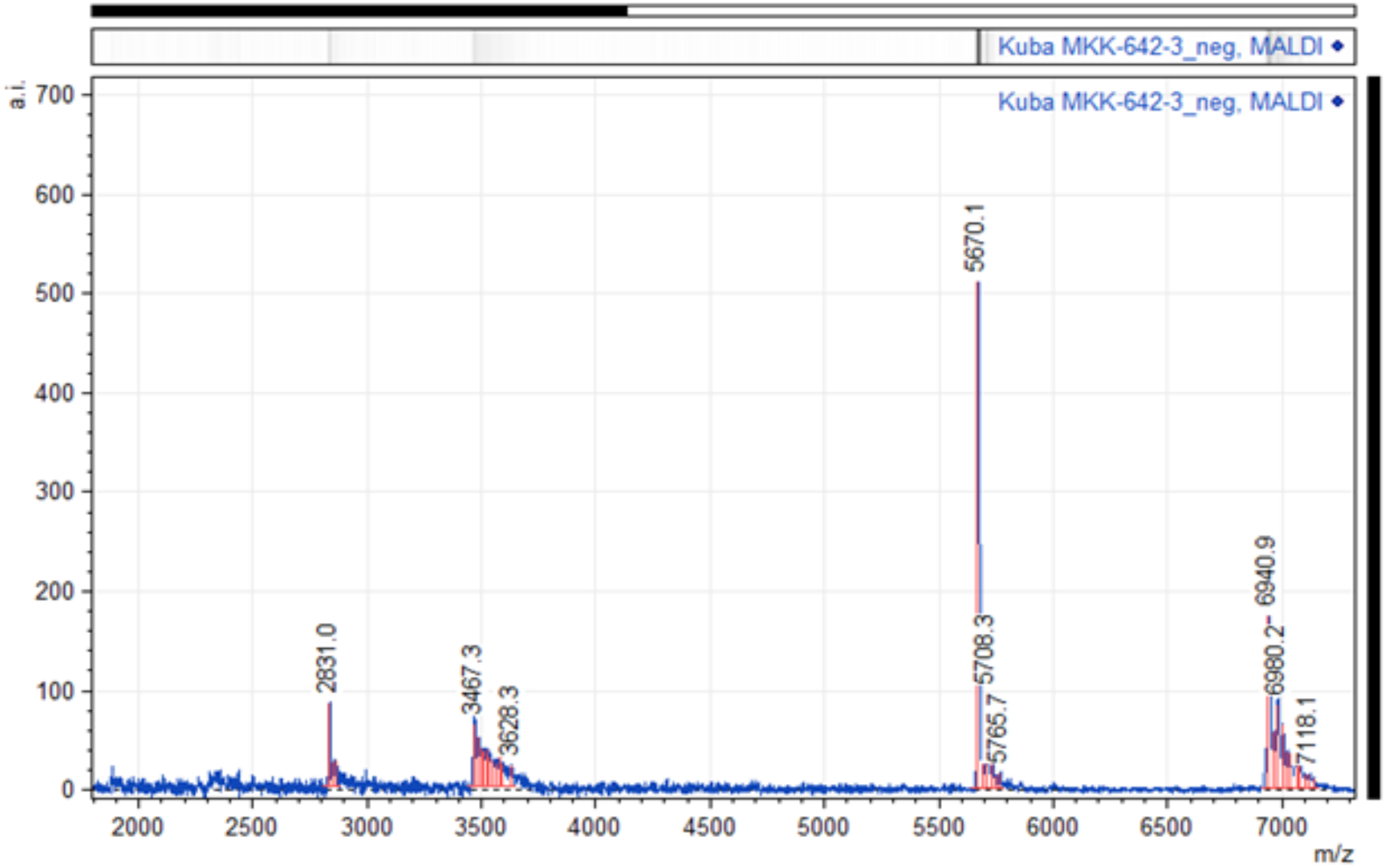
MALDI-TOF mass spectrometry spectrum of single-modified PEX product (ON16_1CSiR) obtained using dCSiRTP. Calculated for [M]: 6942.2 Da; found: 6940.9 Da. The peak at m/z = 5670.1 Da represents template Temp1PEX.

**Figure S8.**
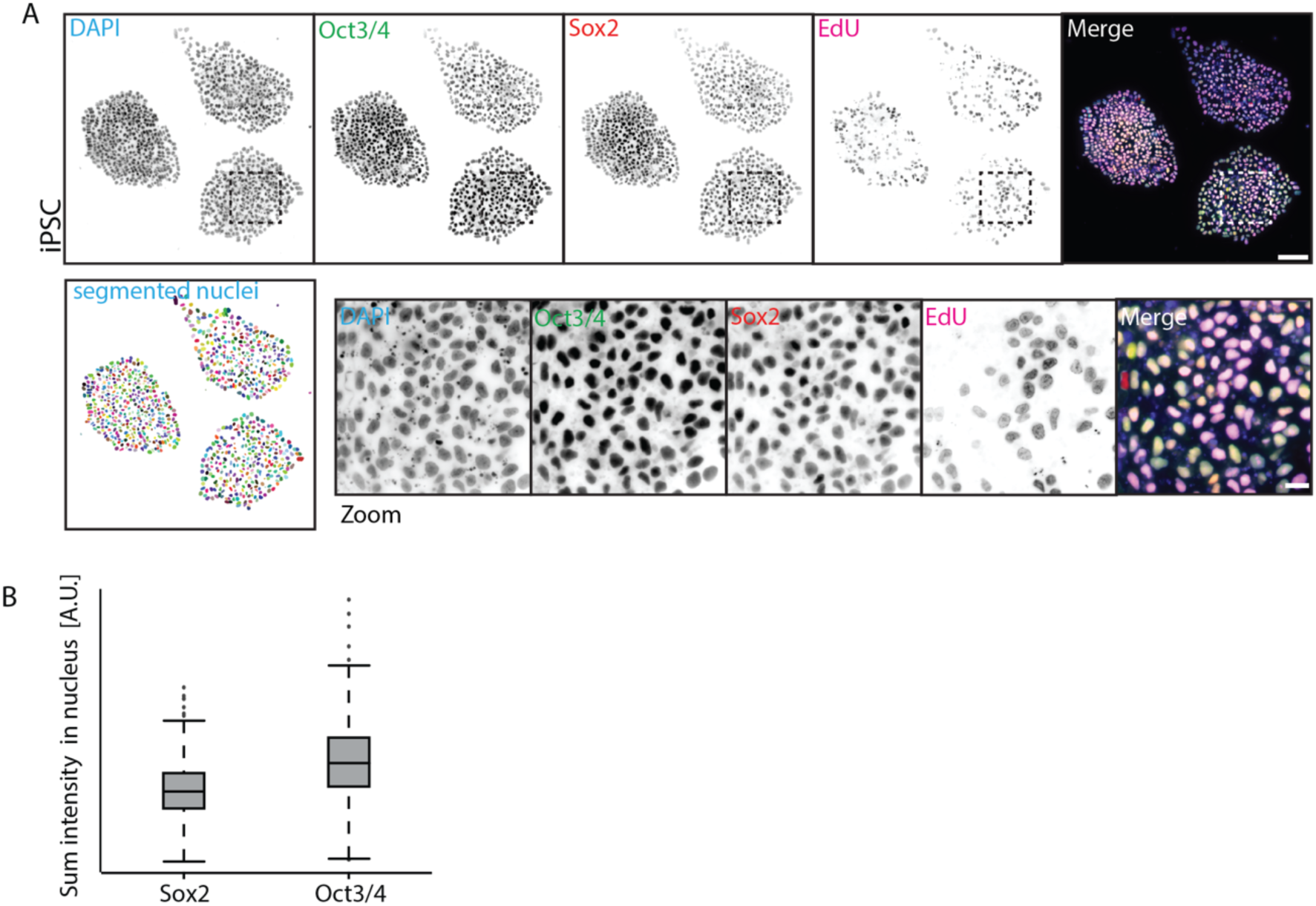
**(A)** Immunofluorescence detection and quantification of pluripotent markers (Oct3/4 and Sox2) and detection of replicating cells using EdU (10 minutes, 10 µm) nucleoside pulse in iPSC cells (Methods). **(B)** Quantification of sum nuclear intensity of Sox2 and Oct3/4 quantified and represented as box plots. Scale bars: 50 µm.

**Figure S9.**
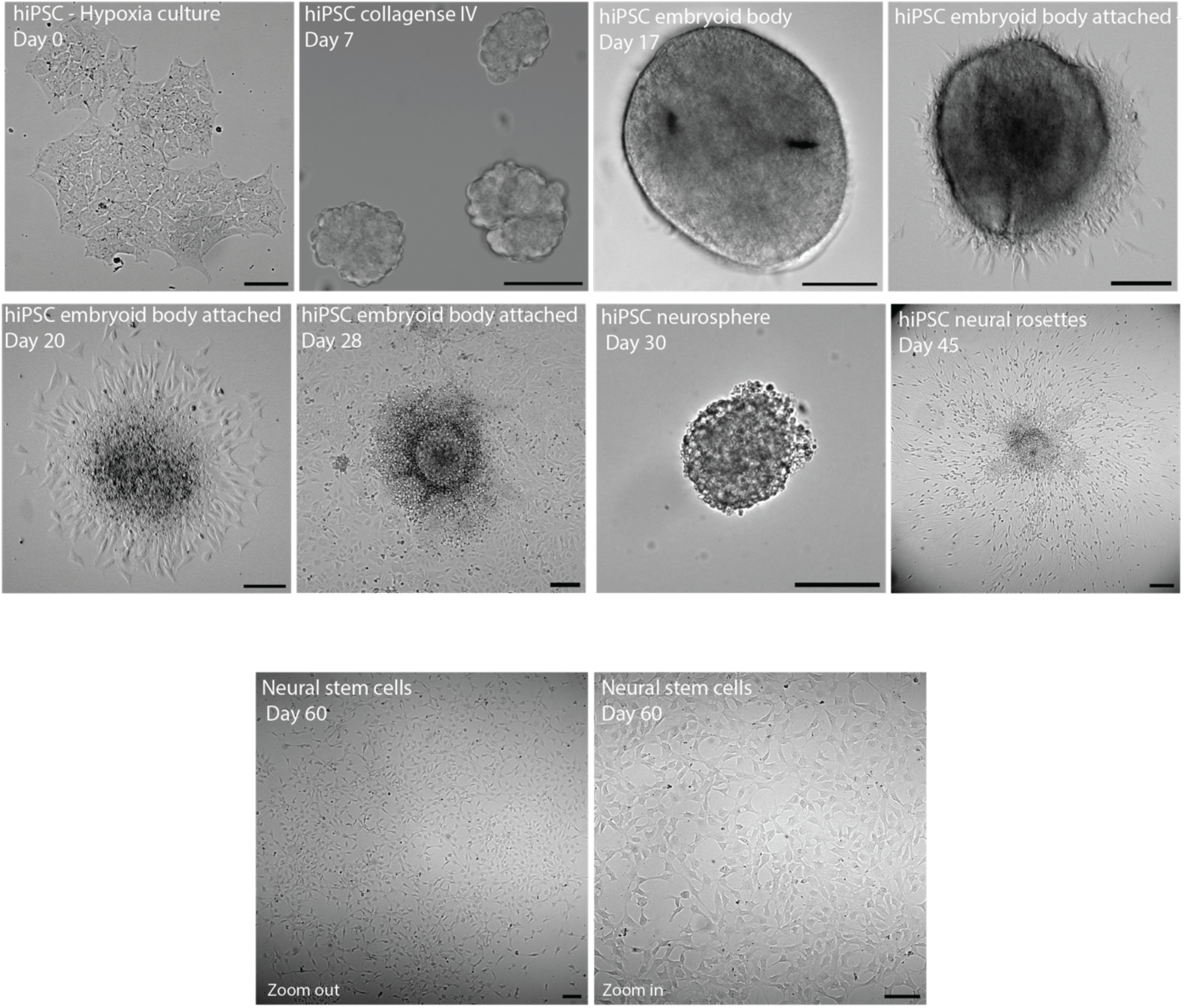
Representative images using differential interference contrast (DIC) microscopy of live cells during the generation of neural stem cells (NSCs) from induced pluripotent stem cells (iPSCs). Scale bars: 100 µm.

**Figure S10.**
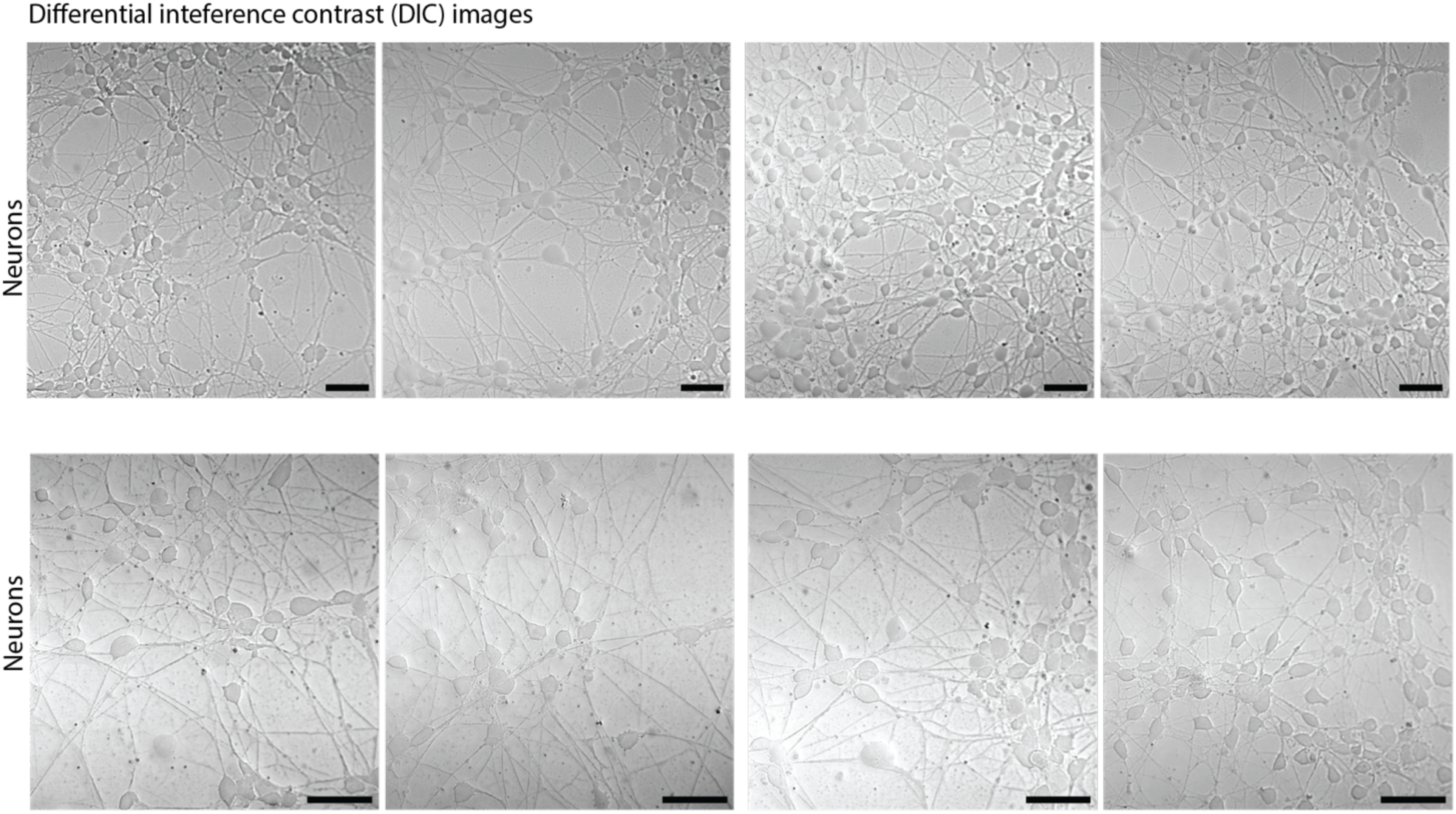
Differential interference contrast (DIC) representative images of differentiated neurons. Scale bars: 25 µm.

**Figure S11.**
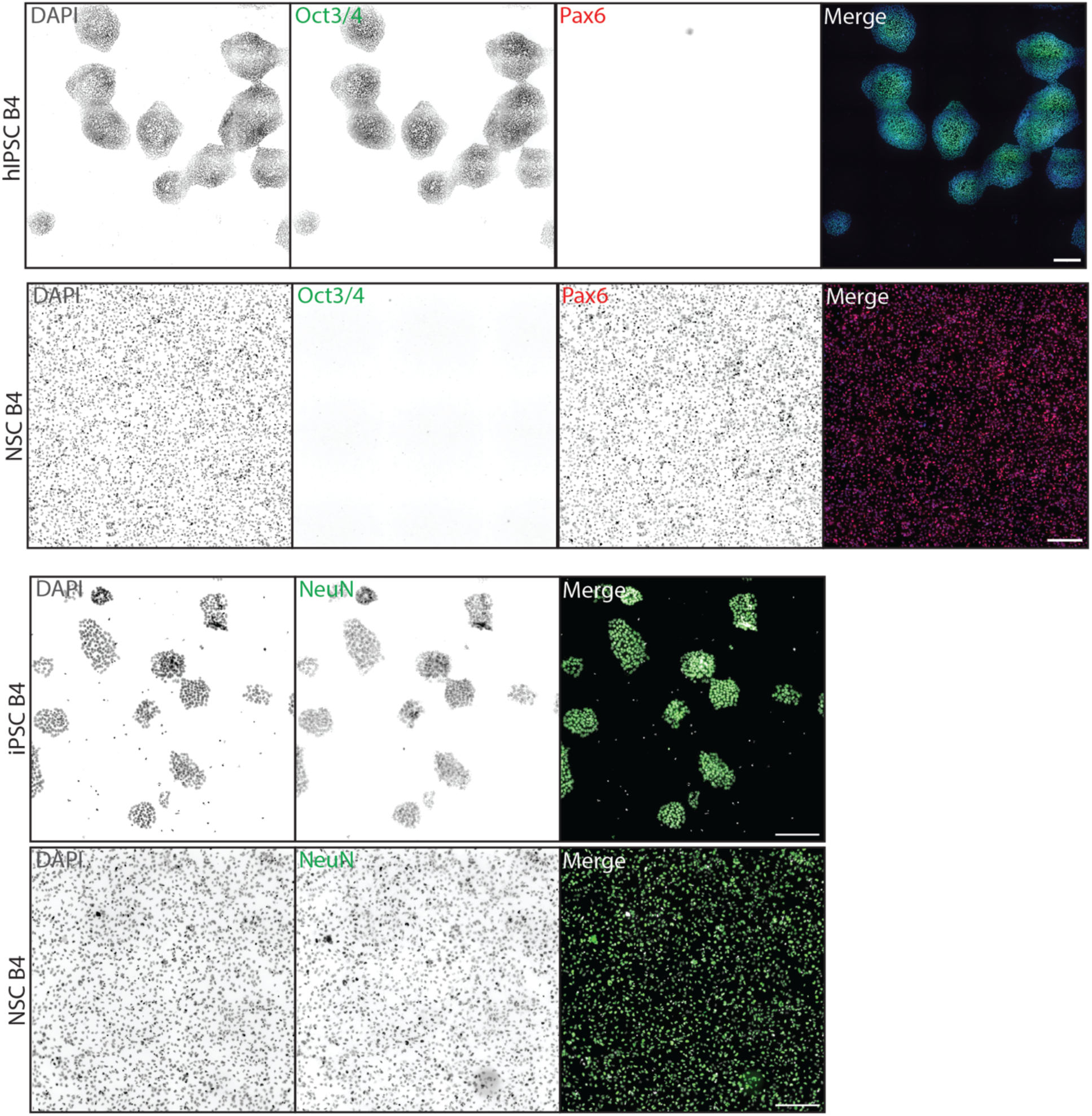
Immunofluorescence detection of pluripotent and neural stem cell markers on IPSCs and NSCs (Table S8, Methods: Immunofluorescence). Scale bars: 250 µm.

**Figure S12.**
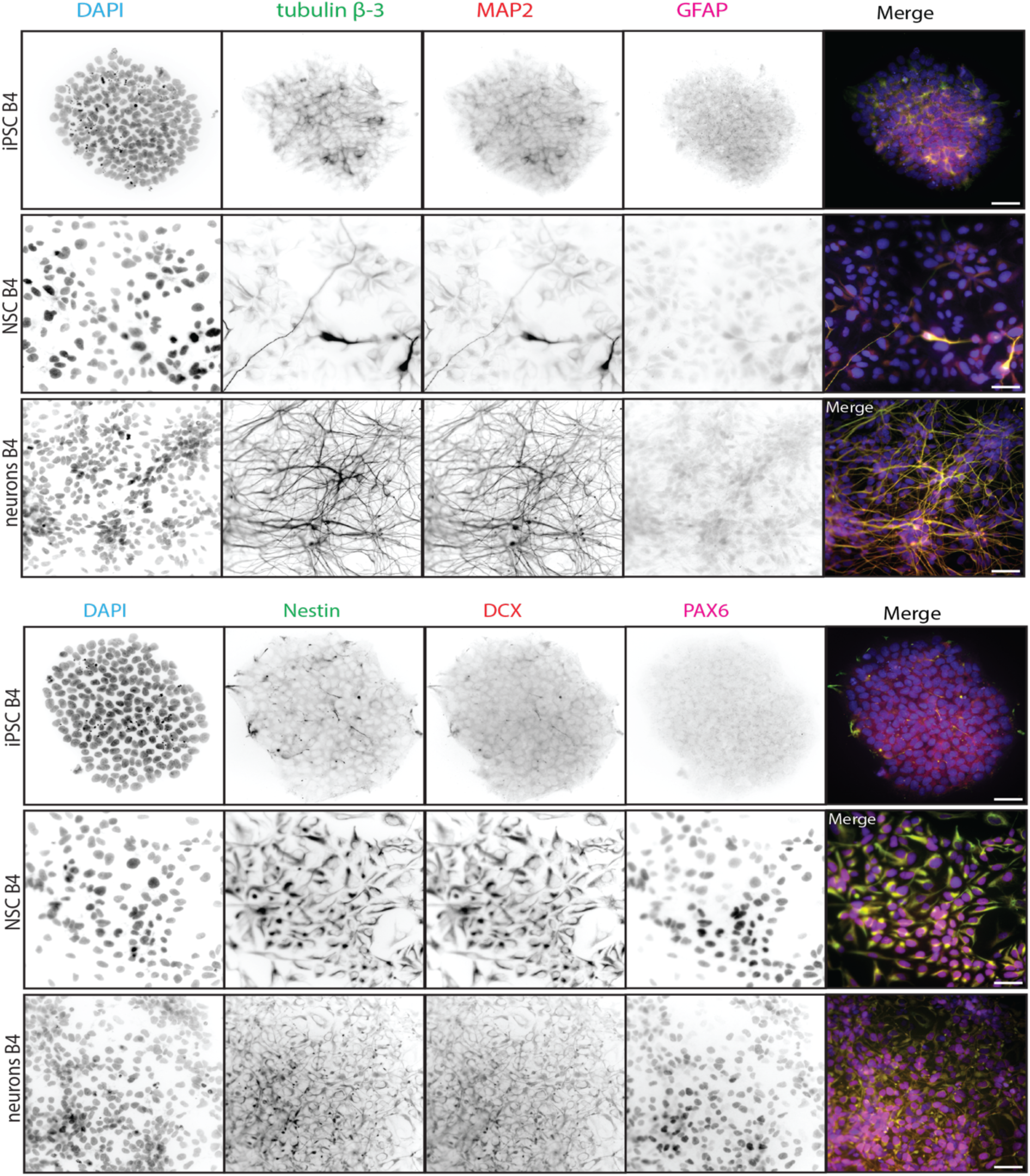
Immunofluorescence detection of neuronal markers on IPSCs, NSCs, and neurons (Table S8, Methods). Scale bars: 50 µm.

**Table S1.**
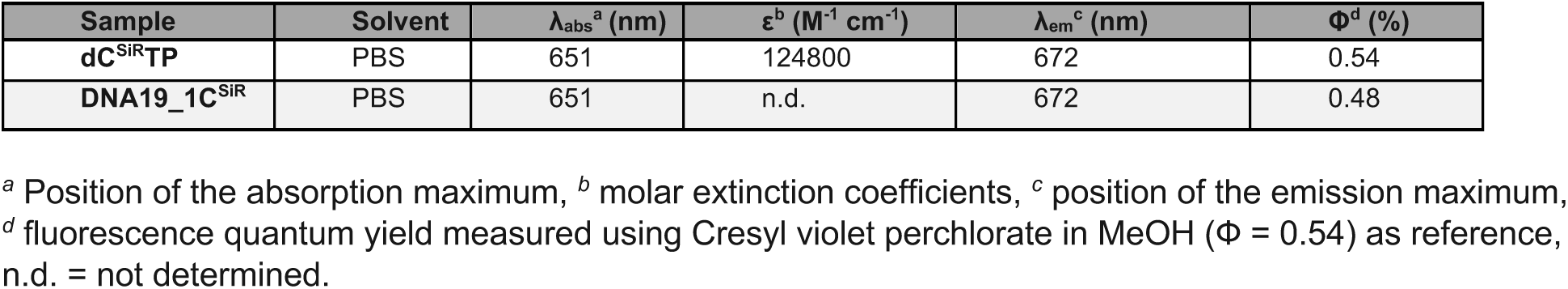
Photophysical properties of dC^SiR^TP and DNA19_1C^SiR^.

| Sample | Solvent | $\lambda_{abs}^a$ (nm) | $\varepsilon^b$ (M<sup>-1</sup> cm<sup>-1</sup>) | $\lambda_{em}^c$ (nm) | $\Phi^d$ (%) |
| --- | --- | --- | --- | --- | --- |
| dC<sup>SiR</sup>TP | PBS | 651 | 124800 | 672 | 0.54 |
| DNA19_1C<sup>SiR</sup> | PBS | 651 | n.d. | 672 | 0.48 |

<sup>a</sup> Position of the absorption maximum, <sup>b</sup> molar extinction coefficients, <sup>c</sup> position of the emission maximum, <sup>d</sup> fluorescence quantum yield measured using Cresyl violet perchlorate in MeOH ( $\Phi = 0.54$ ) as reference, n.d. = not determined.

**Table S2.**
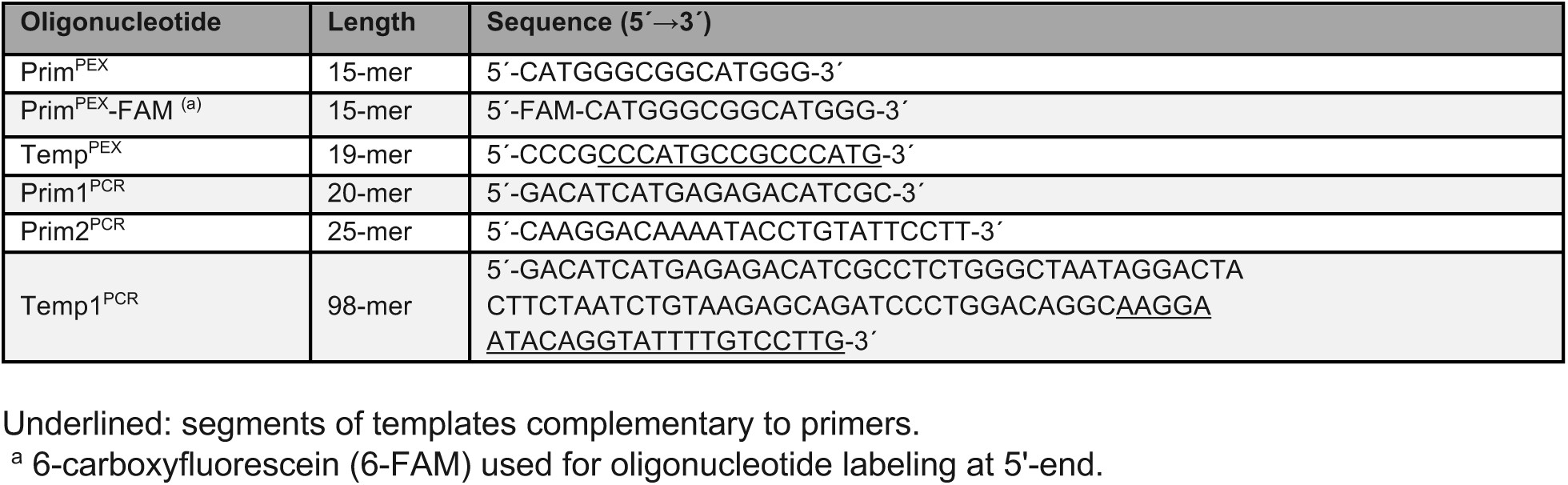
Table of oligonucleotide sequences used in enzymatic synthesis.wi.

**Table S3.**
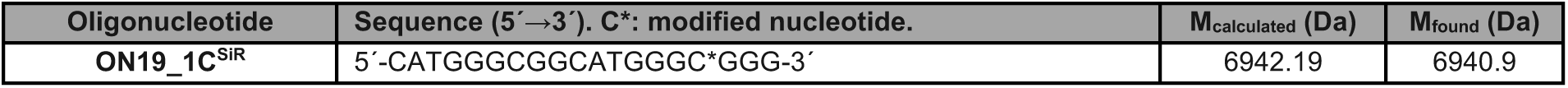
Oligonucleotide prepared, isolated and characterized by MALDI-TOF mass spectrometry.

**Table S4.**
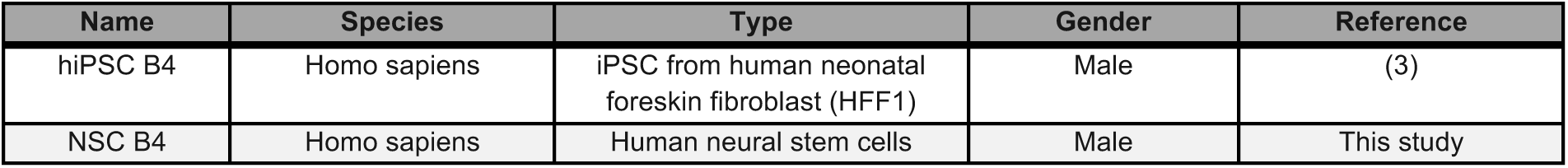
Cell line characteristics.

| Name | Species | Type | Gender | Reference |
| --- | --- | --- | --- | --- |
| hiPSC B4 | Homo sapiens | iPSC from human neonatal foreskin fibroblast (HFF1) | Male | (3) |
| NSC B4 | Homo sapiens | Human neural stem cells | Male | This study |

**Table S5.** Nucleotide and chemical characteristics.

| Name | Application | Detection | Cat # | Company |
| --- | --- | --- | --- | --- |
| dC<sup>SiR</sup>TP | Replication labeling (Labeling of nascent DNA) | - | - | This study |
| NTP-Transporter Live Cell Dye (SNTT 1) | Transport of dC<sup>SiR</sup>TP across cell membrane of living cells | - | SCT064 | Merck, Germany |
| EdU (5-ethynyl-2'-deoxyuridine) | Replication labeling (Labeling of nascent DNA) | Click-IT | 7845.1 | Carl Roth, Germany |
| Abberior DNA live 590 dye | Live cell labeling of total DNA | - | LV 590-0143 | Abberior, Germany |
| Abberior Tubulin live 610 dye | Live cell labeling of tubulin | - | LV 610-0141 | Abberior, Germany |

**Table S6.** Imaging systems characteristics.

| Microscope / Company | Lasers/lamps | Filters (ex. & em. [nm])* | Objectives/ lenses | Detection system | Incubation system | Application |
| --- | --- | --- | --- | --- | --- | --- |
| Zeiss LSM 900 Airyscan, Zeiss, Germany | Four LEDs - (LED-Modul 385 nm, LED-Modul 470nm, LED-Modul 540-580 nm, LED-Modul 625 nm)<br>Laser - Power<br>405 nm- 5 mW<br>488 nm- 10 mW<br>561 nm- 10 mW<br>640 nm- 5 mW | ex: 390/40<br>em: 450/40 &<br>ex: 470/40<br>em: 525/50 &<br>ex: 550/25<br>em: 605/70 &<br>ex: 640/30<br>em: 690/50<br>Quad BP<br>425/30 +<br>514/30 +<br>592/25 +<br>709/100 | EC Plan-Neofluar 2.5x<br>Plan-Apochromat 20x<br>Plan-Apochromat 40x<br>C Plan-Apochromat 63x<br>alpha Plan-Apochromat 100x | **Axiocam 820** mono SONY IMX541 CMOS back illuminated pixel size 2.74 µm x 2.74 µm 4512 x 4512 pixel QE: 86 % at 520 nm Full well capacity 10 000 e 20 fps full frame noise 2.3e at full well capacity **GaAsP Detectors** Additionally an **Airyscan detector** is available. | live cell incubation chamber for temperature, local humidity and CO<sub>2</sub> are installed<br>Definite focus system for hardware focus stabilization<br>AI sample finder for automatic detection of the sample carrier, sample layout and sub-samples<br>photo manipulation is available through the regular user interface | Super resolution microscopy |
| Nikon TiE2 inverted with crest spinning disk unit/ Nikon, Japan | SPECTRA X light engine<br>395/25 nm with 295 mW<br>440/20 nm with 256 mW<br>470/24 nm with 196 mW<br>510/25 nm with 62 mW<br>540/30 nm with 231 mW<br>550/15 nm with 260 mW<br>575/25 nm with 310 mW | LED-DA/FI/TR/Cy5 -4X-B<br>Quadbandpas sex:390/18, 475/35, 535/50<br>em:460/60, 530/43, 580LP | 40x air (0.95 NA) & 250 µm WD*** | Cooled Nikon Qi2 camera and 16.25-megapixel sCMOS sensor. readout noise is: 2.2. electron | Self-build - 37°C incubation chamber, with 5% CO<sub>2</sub> and 60% humidity chamber | high throughput, high content imaging and image analysis |
| STED 775 QUAD Scan microscope (Abberior Instruments) | excitation laser pulsed diode laser 594 nm, <500mW (PDL 594, Abberior Instruments) pulsed diode laser 638nm, 20mW (PiL063X, Advanced Laser Diode Systems) | STED Laser pulsed fiber laser 775 nm, 1200mW (PFL-P-30-775B1R, MPB Communications) 3D Module SLM based (easy 3D, Abberior Instruments) | 100 NA 1.4 Olympus UPlanSApo | Detectors: APD: 605 nm - 625 nm APD: 650nm - 720nm | - | Super resolution microscopy |
| Amersham AI600 imager | Chemiluminescence, UV transillumination | - | - | - | - | Western blots and DNA agarose gels |

\* ex.: excitation & em.: emission, \*\* dichroic specification, \*\*\* WD: working distance.

**Table S7.** Media composition.

| Name | Working solution | Cat # | Company |
| --- | --- | --- | --- |
| IMDM (Iscove's Modified Dulbecco's Medium) | 75 % | 12440053 | Gibco, United States of America |
| Ham's F12 nut mix | 25 % | 21765-029 | Gibco, United States of America |
| penicillin/streptomycin (100x) | 1 x | P0781 | Merck Millipore, Germany |
| N-2 supplement | 0.5 x | N2-K | Capricorn Scientific, Germany |
| B-27™ supplement minus vitamin A | 0.5 x | 12587010 | Thermo Fisher Scientific, United States of America |
| knockout serum replacement | 0.05 % | 10828 | Gibco, United States of America |
| L-ascorbic acid | 50 µg/mL | A5960 | Sigma Aldrich, Germany |
| MTG (1-thioglycerol) | 4 x 10<sup>-4</sup> M | M6145 | Sigma Aldrich, Germany |
| Recombinant Human BMP-1A/ALK-3 Fc Chimera | 500 ng/mL | 315-BR-100/CF | PeproTech, Germany |
| StemMACS™ SB431542 selective inhibitor of ALK5/TGF-β type I receptor) | 10 µM | SB431542 | Miltenyi Biotec, Germany |
| human-FGF2 | 10 ng/mL | 100-18G-250UG | PeproTech, Germany |

*ITSF media*
| Name | Working solution | Cat # | Company |
| --- | --- | --- | --- |
| DMEM / Ham's F12 nutrient mix | 100 % | 10565-018 | Gibco, United States of America |
| Apo-transferrin | 100 µg/mL | 616395 | Sigma Aldrich, Germany |
| Insulin | 25 µg/mL | I6634 | Sigma Aldrich, Germany |
| Fibronectin | 2.5 µg/mL | 33010018 | Thermo Fisher, United States of America |
| Sodium Selenite | 5 ng/mL | S5261 | Sigma Aldrich, Germany |
| penicillin/streptomycin (100x) | 1 x | P0781 | Merck Millipore, Germany |

*Neurosphere media*
| Name | Working solution | Cat # | Company |
| --- | --- | --- | --- |
| DMEM | 33.33 % | 41965039 | Gibco, United States of America |
| DMEM/ Ham's F12 Nut mix | 33.33 % | 10565-018 | Gibco, United States of America |
| Ham's F12 Nut mix | 33.33 % | 21765-029 | Gibco, United States of America |
| N-2 supplement | 0.5 x | N2-K | Capricorn Scientific, Germany |
| penicillin/streptomycin (100x) | 1 x | P0781 | Merck Millipore, Germany |
| h-FGF2 | 10 ng/mL | 100-18G-250UG | PeproTech, Germany |

*NSC media*
| Name | Working solution | Cat # | Company |
| --- | --- | --- | --- |
| DMEM | 33.33 % | 41965039 | Gibco, United States of America |
| DMEM/ Ham's F12 Nut mix | 33.33 % | 10565-018 | Gibco, United States of America |
| Ham's F12 Nut mix | 33.33 % | 21765-029 | Gibco, United States of America |
| N-2 supplement | 1 x | N2-K | Capricorn Scientific, Germany |
| B-27™ supplement minus vitamin A | 0.5 x | 12587010 | Thermo Fisher Scientific, United States of America |
| Insulin | 20 µg/mL | I6634 | Sigma Aldrich, Germany |
| penicillin/streptomycin (100x) | 1 x | P0781 | Merck Millipore, Germany |
| human-EGF | 10 ng/mL | 100-15 | PeproTech, Germany |
| human-FGF2 | 10 ng/mL | 100-18G-250UG | PeproTech, Germany |

Differentiation media
| Name | Working solution | Cat # | Company |
| --- | --- | --- | --- |
| Neurobasal media | - | 21103049 | Thermo Fisher Scientific, United States of America |
| B-27™ supplement minus vitamin A | 1 x | 12587010 | Thermo Fisher Scientific, United States of America |
| penicillin/streptomycin (100x) | 1 x | P0781 | Merck Millipore, Germany |
| Glutamax | 1 x | 35050038 | Thermo Fisher Scientific, United States of America |

**Table S8.**
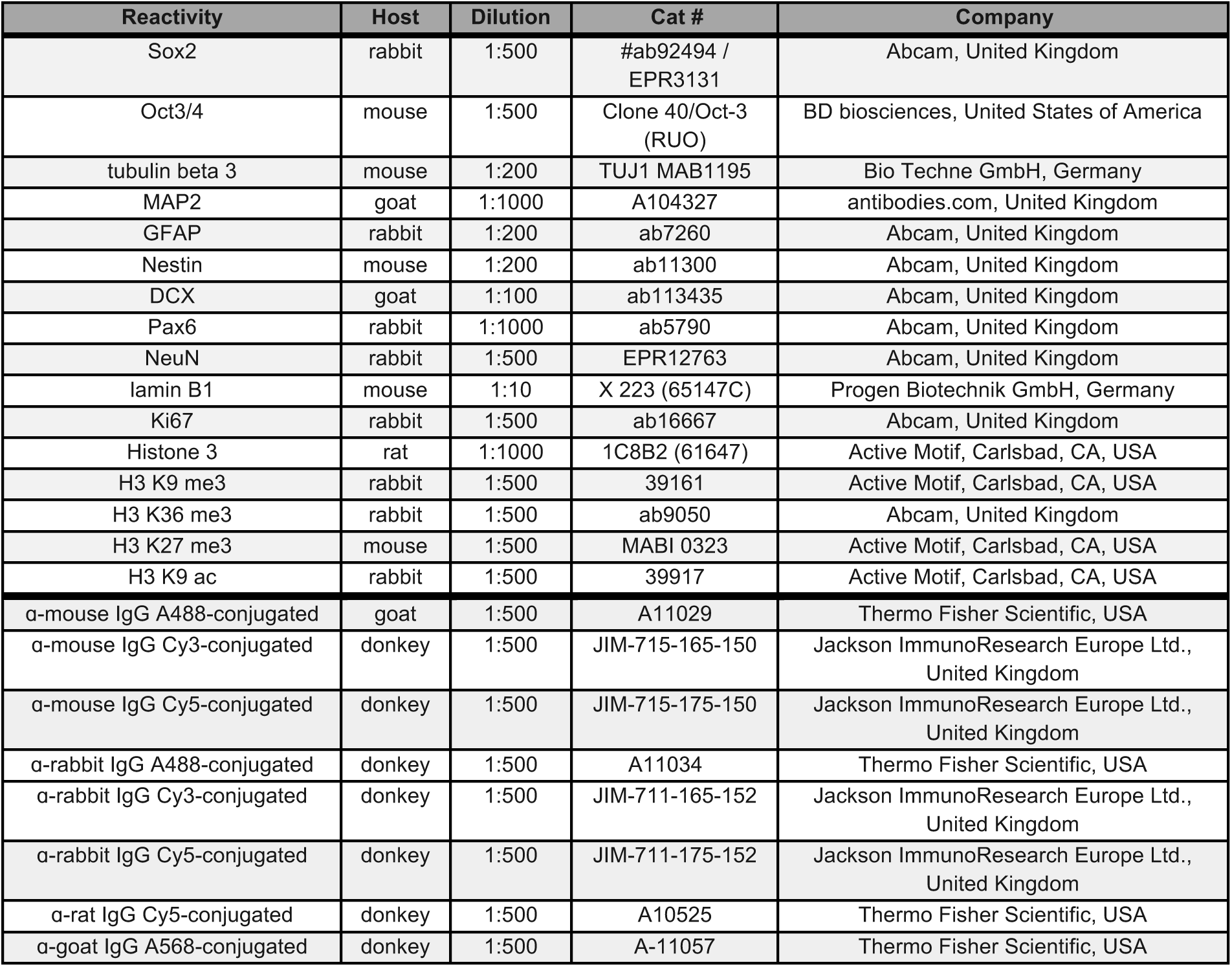
Primary and secondary antibody characteristics.

| Reactivity | Host | Dilution | Cat # | Company |
| --- | --- | --- | --- | --- |
| Sox2 | rabbit | 1:500 | #ab92494 / EPR3131 | Abcam, United Kingdom |
| Oct3/4 | mouse | 1:500 | Clone 40/Oct-3 (RUO) | BD biosciences, United States of America |
| tubulin beta 3 | mouse | 1:200 | TUJ1 MAB1195 | Bio Techne GmbH, Germany |
| MAP2 | goat | 1:1000 | A104327 | antibodies.com, United Kingdom |
| GFAP | rabbit | 1:200 | ab7260 | Abcam, United Kingdom |
| Nestin | mouse | 1:200 | ab11300 | Abcam, United Kingdom |
| DCX | goat | 1:100 | ab113435 | Abcam, United Kingdom |
| Pax6 | rabbit | 1:1000 | ab5790 | Abcam, United Kingdom |
| NeuN | rabbit | 1:500 | EPR12763 | Abcam, United Kingdom |
| lamin B1 | mouse | 1:10 | X 223 (65147C) | Progen Biotechnik GmbH, Germany |
| Ki67 | rabbit | 1:500 | ab16667 | Abcam, United Kingdom |
| Histone 3 | rat | 1:1000 | 1C8B2 (61647) | Active Motif, Carlsbad, CA, USA |
| H3 K9 me3 | rabbit | 1:500 | 39161 | Active Motif, Carlsbad, CA, USA |
| H3 K36 me3 | rabbit | 1:500 | ab9050 | Abcam, United Kingdom |
| H3 K27 me3 | mouse | 1:500 | MABI 0323 | Active Motif, Carlsbad, CA, USA |
| H3 K9 ac | rabbit | 1:500 | 39917 | Active Motif, Carlsbad, CA, USA |
| α-mouse IgG A488-conjugated | goat | 1:500 | A11029 | Thermo Fisher Scientific, USA |
| α-mouse IgG Cy3-conjugated | donkey | 1:500 | JIM-715-165-150 | Jackson ImmunoResearch Europe Ltd., United Kingdom |
| α-mouse IgG Cy5-conjugated | donkey | 1:500 | JIM-715-175-150 | Jackson ImmunoResearch Europe Ltd., United Kingdom |
| α-rabbit IgG A488-conjugated | donkey | 1:500 | A11034 | Thermo Fisher Scientific, USA |
| α-rabbit IgG Cy3-conjugated | donkey | 1:500 | JIM-711-165-152 | Jackson ImmunoResearch Europe Ltd., United Kingdom |
| α-rabbit IgG Cy5-conjugated | donkey | 1:500 | JIM-711-175-152 | Jackson ImmunoResearch Europe Ltd., United Kingdom |
| α-rat IgG Cy5-conjugated | donkey | 1:500 | A10525 | Thermo Fisher Scientific, USA |
| α-goat IgG A568-conjugated | donkey | 1:500 | A-11057 | Thermo Fisher Scientific, USA |

**Table S9.**
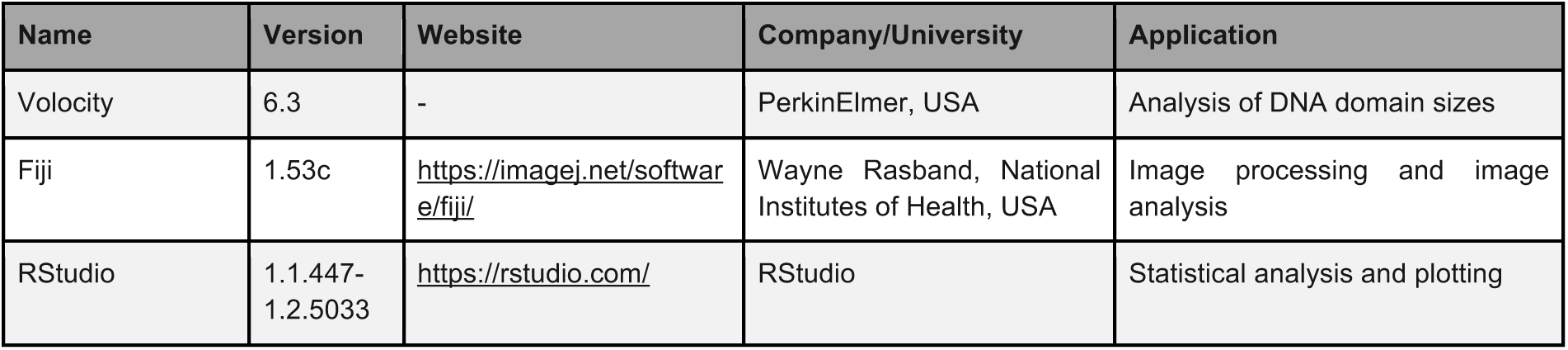
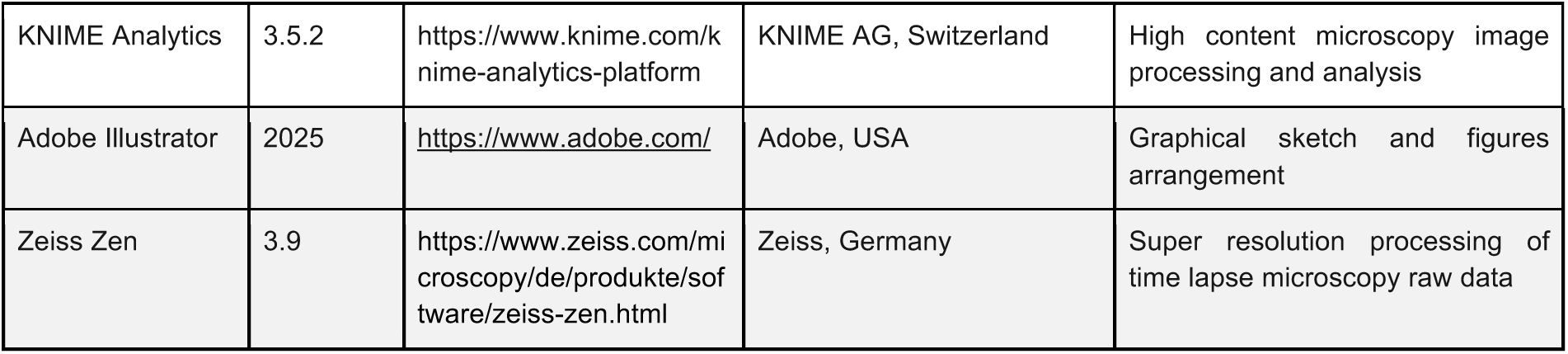
Software.

| Name | Version | Website | Company/University | Application |
| --- | --- | --- | --- | --- |
| Volocity | 6.3 | - | PerkinElmer, USA | Analysis of DNA domain sizes |
| Fiji | 1.53c | https://imagej.net/software/fiji/ | Wayne Rasband, National Institutes of Health, USA | Image processing and image analysis |
| RStudio | 1.1.447-1.2.5033 | https://rstudio.com/ | RStudio | Statistical analysis and plotting |
| KNIME Analytics | 3.5.2 | https://www.knime.com/knime-analytics-platform | KNIME AG, Switzerland | High content microscopy image processing and analysis |
| Adobe Illustrator | 2025 | https://www.adobe.com/ | Adobe, USA | Graphical sketch and figures arrangement |
| Zeiss Zen | 3.9 | https://www.zeiss.com/microscopy/de/produkte/software/zeiss-zen.html | Zeiss, Germany | Super resolution processing of time lapse microscopy raw data |

